# Active enhancers strengthen insulation by RNA-mediated CTCF binding at TAD boundaries

**DOI:** 10.1101/2021.07.13.452118

**Authors:** Zubairul Islam, Bharath Saravanan, Kaivalya Walavalkar, Jitendra Thakur, Umer Farooq, Anurag Kumar Singh, Radhakrishnan Sabarinathan, Awadhesh Pandit, Steven Henikoff, Dimple Notani

## Abstract

Vertebrate genomes are partitioned into Topologically Associating Domains (TADs), which are typically bound by head-to-head pairs of CTCF binding sites. Transcription at TAD boundaries correlates with better insulation, however, it is not known whether the boundary transcripts themselves contribute to boundary function. Here we characterize boundary-associated RNAs genome-wide, focusing on the disease-relevant *INK4a/ARF* TAD. Using CTCF site deletions and boundary-associated RNA knockdowns, we observe that boundary-associated RNAs facilitate recruitment and clustering of CTCF at TAD borders. The resulting CTCF enrichment enhances TAD insulation, enhancer:promoter interactions and TAD gene expression. Importantly, knockdown of boundary-associated RNAs results in loss of boundary insulation function. Using enhancer deletions and CRISPRi of promoters we show that active TAD enhancers but not promoters induce boundary-associated RNA transcription, thus defining a novel class of regulatory enhancer RNAs.

## Introduction

The genomes of multicellular eukaryotes are organized into self-interacting units known as Topologically Associating Domains (TADs) (1). TADs are identified in Hi-C interaction matrices as contact domains, and contact domains that exhibit corner dots (at the apex of a loop) are known as loop domains (2). Loop and contact domains are both proposed to be formed by loop extrusion (1,3-7) and are dependent on CTCF and cohesin (6-10). The length of an extruded loop is determined by the presence and orientation of CTCF bound regions that interact within the loop (11). By preventing enhancers and promoters within the loop from interacting outside of the loops, boundary CTCF pairs insulate genes within a TAD (7-8,12-13). However, the presence of CTCF consensus motifs or their orientation at contact domains do not accurately predict TAD boundaries, suggesting that additional factors must influence CTCF binding and, therefore, TAD formation and insulation.

Mutation of CTCF sites at TAD boundaries affects gene expression, ranging from no effect to severe effect in the same and neighboring TADs (14-21). Mutation of all CTCF sites within a boundary region or depletion of CTCF may be required to display effects on transcription (20,22), suggesting redundancy among multiple CTCF sites at a TAD boundary. However, it is unclear how multiple CTCF sites within the boundary coordinate to achieve better insulation. Active transcription at TAD boundaries correlates with better insulation and CTCF occupancy (22-24) and has been speculated to increase TAD strength (25-27), consistent with observations that inhibition of transcription or Ribonuclease A (RNaseA) treatment disrupts 3D chromatin architecture (28-30). Furthermore, CTCF dimerization, important for loop extrusion, is RNA-dependent, and RNA binding of CTCF is required for TAD chromatin organization (31-32), but how CTCF:RNA interactions maintain TADs is not clear.

In some instances, TAD boundaries are found near sites of enhancers (33). Also, TADs containing super-enhancers consist of multiple enhancers exhibit strong boundaries (25,34). However, it is not known whether these enhancers play a role in boundary insulation. Moreover, boundaries themselves display enhancer-like features (30,35), although the levels of H3K27ac, RNA Polymerase II (PolII) and H3K4me3 on TAD boundaries are far less than are found on active enhancers in general (36). These observations suggest a relationship between certain TAD boundaries and enhancers, but the mechanistic basis for their putative interactions remains unknown.

Here we show that boundaries of about half of human TADs are transcribed, either from a gene-body that overlaps a boundary region or from a non-genic region that is transcribed within the boundary. Compared to non-transcribed TAD boundaries, transcribed genic and non-genic boundaries are more enriched for CTCF binding and are better insulators of genes within TADs, which are more highly expressed. RNAs transcribed from non-genic (i.e. enhancer-like regions) boundaries are more stable than RNAs expressed from enhancers at non-boundary regions. Similarly, genic transcripts from boundaries are also more stable than other mRNAs. To test for functional relevance of boundary-associated RNAs, we focused on the well-characterized *INK4a/ARF* TAD, which is bound by CTCF sites and harbors a super-enhancer. Both deletions of CTCF sites and knockdown of boundary-associated RNAs resulted in loss of enhancer:promoter interactions within TADs while strengthening interactions with enhancers in regions outside the TAD. These observations suggest that active enhancers within TADs promote TAD insulation by triggering transcription of the boundaries. We propose that the RNAs produced at boundaries recruit CTCF, resulting in robust intra-TAD transcription.

## Results

### Transcribed TAD boundaries insulate better than non-transcribed boundaries

To determine the frequency of transcribed TAD boundaries, we measured run-on transcription with GRO-seq (37) on TAD boundaries identified by HOMER’s findTADsAndLoops.pl. This program identifies both contact and loop domain TADs based on intra-domain interaction frequency. Both TADs and loops are thought to be formed by loop extrusion and to be dependent on CTCF and cohesin (6, 8-10). HOMER identified 15,397 TAD boundaries at 5 kb resolution using HiC data from HeLa cells (11). GRO-seq detected at least ten normalised counts within about half of the TADs identified by HOMER (8134, 52.8%, Fig. 1A). Almost 80% of transcribed boundaries are genic (Fig. 1B-C).

**Figure 1.**
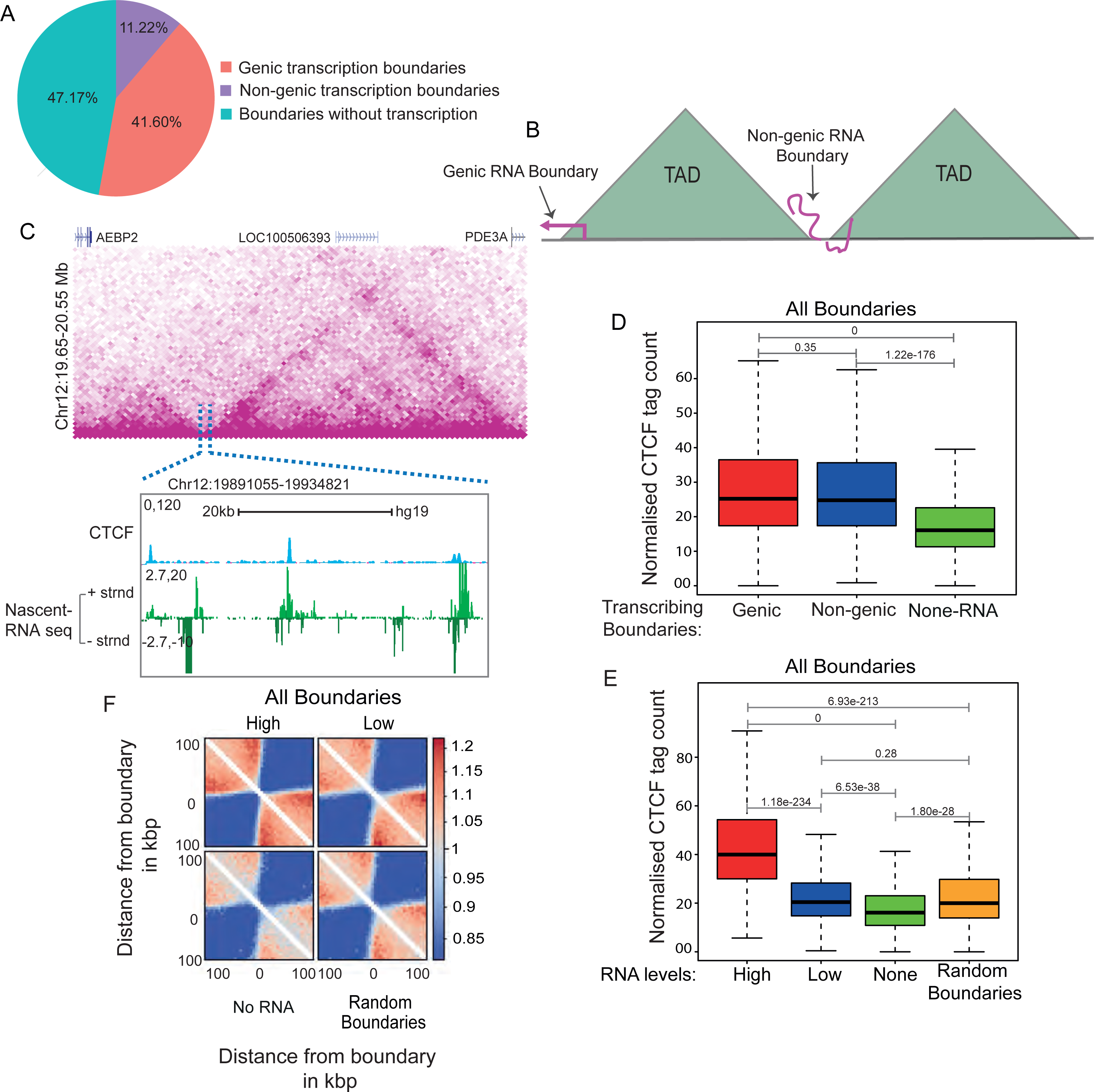
Transcribed TAD boundaries insulate better than non-transcribed boundaries: **A.** Pie chart displaying the percentage of boundaries that show genic transcription, non-genic de novo transcription and no transcription. **B.** Schematic depicting genic and non-genic transcribed boundaries. **C.** Browser shot displaying a TAD structure from HiC data. The zoomed in box shows a *de novo* non-genic transcribed boundary region that is overlaid with CTCF and mNET-seq signal. **D.** Box plots showing CTCF enrichment on genic, non-genic and non-transcribed boundaries. **E.** Box plots showing CTCF enrichment on all boundaries exhibiting varying levels of RNAs versus non-transcribed and random boundaries. **F.** Pile up (Aggregated normalized HiC interactions) plot centered at high transcribed, low transcribed, non-transcribed and random boundaries at a 5 kb resolution. The 100 kb distances are taken from the boundary region. The p-values in boxplots were calculated using the Wilcoxon rank-sum test. The boxplots depict the minimum (Q1-1.5*IQR), first quartile, median, third quartile and maximum (Q3+1.5*IQR) without outliers.

TAD boundaries are enriched for CTCF binding. We determined if strong CTCF binding at TAD boundaries is associated with increased transcription. Using high-resolution CTCF CUT&RUN data (38), we observed that both genic and non-genic transcribed TAD boundaries had significantly higher CTCF enrichment than non-transcribed boundaries (Fig. 1D). This relationship between transcription and CTCF enrichment at TAD boundaries was also observed for different RNA levels (Fig. 1E). Within non-genic transcribed boundaries, CTCF enrichment was also correlated with GRO-seq signal (Fig. S1A-B). Similar results were seen using CTCF ChIP-seq (39) (Fig. S1C-E). We conclude that transcribed TAD boundaries are more enriched for CTCF binding than non-transcribed boundaries.

We next asked if transcribed TAD boundaries that show high CTCF binding are stronger insulators. We noted a positive correlation between the levels of RNA at the transcribed boundaries and the TAD insulation (Fig. 1F). Even TADs with the least transcribed boundaries were better insulated as compared to non-transcribed boundaries (Fig. 1F). We observed a similar positive correlation between boundary RNA levels and TAD insulation using two different bin sizes (5 kb and 20 kb) in our analyses, thereby ruling out resolution-related biases (Fig. S2). A total of 69.2% (10658 out of 15397) of TADs boundaries were found to be transcribed (Fig. S2A). The CTCF enrichment was stronger on transcribed genic and non-genic boundaries at both 5 kb and 20 kb bin sizes (Fig. S2B-E). Insulation scores on transcribed genic and non-genic boundaries were correlated with levels of RNA at the boundaries (Fig. S2F-H)

We also examined the relationship between transcription of TAD boundaries and CTCF binding using HICCUPS, which identifies loop anchors. In total, 3525 loop anchors were identified with 5 kb bin size, of which 43% (1519) were genic and 19% (675) were non-genic (Fig. S3A). As was the case for TAD boundaries identified by HOMER, we found a positive correlation between transcription of loop anchors and CTCF enrichment using both CUT&RUN (Fig. S3B-C) and ChIP-seq datasets (Fig. S3D-E). For subsequent analyses, we focused on TADs called by HOMER with 5 kb boundary regions.

We also observed positive associations between CTCF enrichment, insulation score and transcription at TAD boundaries in IMR90 fibroblasts (11) (Fig. S4A-D) and lymphoblastoid cells K562 (40) (Fig. S4E-H). Together, our results suggest that CTCF enrichment and enhanced insulation are general features of transcribed TAD boundaries.

### RNA increases CTCF occupancy at TAD boundaries

We wondered whether elevated levels of transcription and CTCF at TAD boundaries are causally related. To test this possibility, we treated HeLa cells with RNaseA and performed Western blot analysis on the nuclear and chromatin fractions. We observed a significant loss of CTCF in the chromatin fraction (Fig. 2A, S6A), consistent with the reported dependence of CTCF on RNA for chromatin binding (29,31-32). To map the genomic sites of RNA-dependent CTCF binding, we performed CUT&RUN for CTCF with RNaseA treatment (CUT&RUN.RNaseA). Upon RNA depletion, we observed an overall decrease in CTCF occupancy, especially at the TAD boundaries where CTCF is most enriched (Fig. 2B). Genome-wide, transcribed boundaries exhibited greater loss of CTCF with RNaseA treatment as compared to non-transcribed and randomly selected boundaries (Fig. 2C).

**Figure 2.**
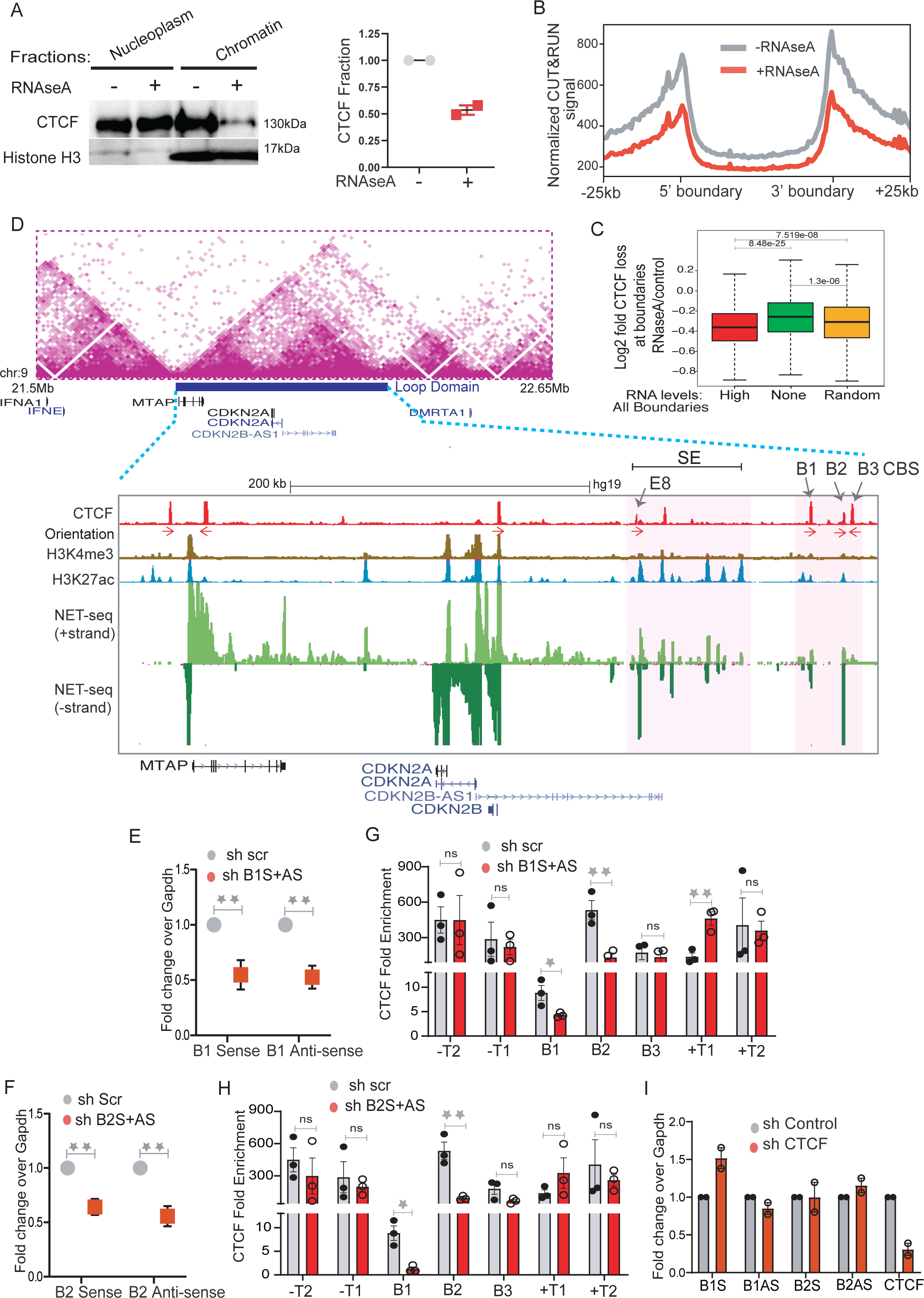
RNA increases CTCF occupancy at TAD boundaries. **A.** Immunoblotting with CTCF and Histone H3 on soluble nucleoplasm and chromatin bound fractions of the nucleus treated with or without RNaseA (Left panel). The right panel shows the quantified loss of CTCF from chromatin fractions from two replicates. **B.** Normalised CTCF CUT&RUN signal in control and RNAseA-treated conditions plotted at equally scaled TADs of 50 kb length each and 25 kb flanks**. C.** Box plot showing the fold loss in CTCF upon RNAseA treatment at boundaries with high RNA levels, non-transcribed and random boundaries. **D.** TAD structure at *INK4a/ARF* locus on 9p21 region (Rao et al., 2014). The loop domain (Blue bar) is overlaid by the position of CTCF peaks, CTCF motif orientation, H3K4me3, H3K27ac (enhancers), mNET-seq tracks and gene annotations. The highlighted region shows the B1, B2 and B3 CTCF sites on the 3’ boundary and super-enhancer with the position of enhancer E8 marked by an arrow. **E.** qRT-PCRs showing the levels of sense and anti-sense non-coding RNA at B1 CTCF site upon B1 RNA knockdown. **F.** qRT-PCRs showing the levels of sense and anti-sense RNA at the B2 CTCF site upon B2 RNA knockdown. **G.** CTCF ChIP enrichment before and after shRNA mediated knockdown of sense and anti-sense RNA at B1 CTCF sites. The bars show CTCF fold enrichment on three CTCF sites (B1, B2 and B3) at the 3’ boundary, on -T1 (5’ boundary) of the *INK4a/ARF* TAD, on -T2 on the adjacent TAD boundary upstream, +T1 (on the adjacent TAD boundary downstream) and +T2, the boundary of the following TAD downstream. **H.** CTCF ChIP enrichment before and after shRNA mediated knockdown of sense and anti-sense RNA at B2 CTCF sites. The bars show CTCF on three CTCF sites (B1, B2 and B3) at the 3’ boundary, on -T1 (5’ boundary) of the *INK4a/ARF* TAD, on -T2 of the adjacent TAD boundary upstream, on +T1 of the adjacent TAD boundary downstream, and on +T2 of the boundary of the following TAD downstream. **I.** qRT-PCRs of sense and anti-sense RNAs from B1 and B2 CTCF sites upon CTCF knockdown. The drops in CTCF levels are shown in the last two bars. Error bars denote SEM from three biological replicates. *p*-values were calculated by Student’s two-tailed unpaired *t*-test in **E, F, G and H.** \*\**p* < 0.01,\**p* < 0.05, ^ns^*p* > 0.05.

We next asked if the loss of CTCF at TAD boundaries is due to the digestion of RNA at direct binding sites as opposed to a general effect of RNA degradation. We chose the ∼500-kb *INK4a/ARF* TAD, which spans the *CDKN2A*, *CDKN2B* and *MTAP* cell cycle regulatory genes and a super-enhancer (Fig. 2D, highlighted region) (41-42). The super-enhancer within this TAD contains several active enhancers that are enriched for H3K27ac modification and enhancer RNAs (eRNAs) (Fig. 2D, mNET-seq panel) (41). We confirmed 5’ TAD boundary locations on the transcribed gene *MTAP* and 3’ boundary on the non-genic region containing three CTCF ChIP-seq peaks (B1, B2 and B3). The position of these boundary regions was persistent in HiC data from other cell types (Data not shown). CTCF peaks B1 and B2 but not B3 exhibit strong nascent transcription based on publicly available mNET-seq, GRO-seq and PRO-seq data (43-45) (Fig. S5) and so are classified as eRNAs. In our CUT&RUN.RNaseA data, we observed a loss of CTCF at the boundary regions of the *INK4a/ARF* TAD (Fig. S6B), which suggests that eRNAs help maintain CTCF at B1 and B2 CTCF sites. To test this interpretation, we depleted sense and anti-sense eRNAs at B1 and B2 CTCF sites using shRNAs (Fig. 2E-F) and confirmed the drop of these eRNAs from chromatin fraction by qRT-PCR (Fig. S6C). Knockdowns using shRNAs to B1 and B2 but not B3 and nearby TAD boundaries (T1 and T2) resulted in the loss of CTCF binding at these sites (Fig. 2G-H and Fig. S6D). Notably, the loss of RNAs at B1 reduced CTCF occupancy not only at B1 but also at B2 and vice versa (Fig. 2G-H). This suggests that CTCF on transcribed boundaries is maintained in part by eRNAs, and different transcribed CTCF sites within a boundary are potentially interdependent. Notably, shRNA-mediated knockdown or over-expression of CTCF had at most minor effects on eRNA expression (Fig. 2I and Fig. S6E), which suggests that CTCF enrichment at TAD boundaries depends on RNA, but not vice-versa.

### Non-coding RNAs and CTCF interact at the *INKa/ARF* TAD boundary

We next asked whether RNA and CTCF interact at functional boundaries. We first performed RNA pulldowns of sense and anti-sense RNAs transcribed from B1 and B2 CTCF sites in the *INKa/ARF* TAD followed by immunoblotting with a CTCF antibody (Fig. 3A). We observed CTCF interactions with B1 and B2 sense and anti-sense RNA and also with an eRNA from the CTCF-bound E8 enhancer within the super-enhancer, but not with the Box B negative control. CTCF binding to B1 and B2 sense and anti-sense RNAs was confirmed by UV-RIP, an RNA immunoprecipitation method (Fig. 3B). Surprisingly, the CTCF motifs present at transcribed boundaries are much sparser than at non-transcribed or randomly selected boundaries (Fig. 3C-D). However, we found that transcribed boundaries are more densely packed with CTCF peaks than are non-transcribed or randomly selected boundaries (Fig. 3E). These data suggest that transcription or the presence of eRNA can reinforce CTCF binding to multiple weak CTCF motifs.

**Figure 3.**
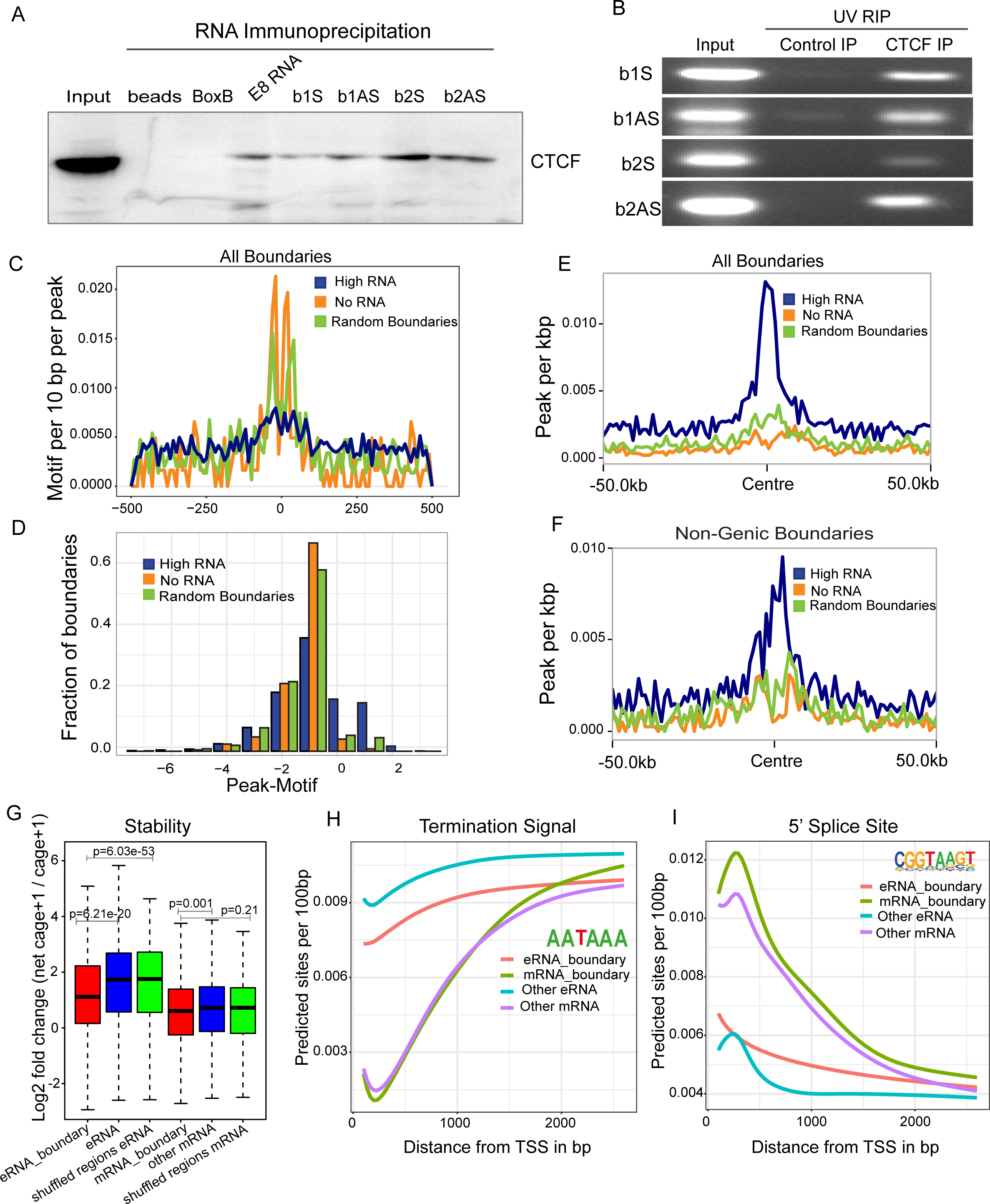
Non-coding RNAs and CTCF interact at the *INKa/ARF* TAD boundary: **A.** Immunoblot for CTCF on RNA pulldowns performed on various RNAs (n=2). **B.** Agarose gel image showing the UV-RIP RT-PCRs for sense and anti-sense RNA arising from B1 and B2 CTCF sites (n=2). **C.** Line plots show the CTCF motif per 10 base pair per peak on various classes of boundaries. **D.** Fraction of boundaries with differential occupancy of CTCF peaks over the presence of the CTCF motif**. E.** CTCF peaks per kb on all transcribed boundaries with high RNA, no RNAs and random boundaries (50 kb flanks on both side). **F.** CTCF peaks per kb on non-genic transcribed boundaries with high RNAs, no RNAs and random boundaries. **G.** Log_2_FC of NET-CAGE+1/CAGE+1, indicating stability of RNA at TSSs identified from NET-CAGE at boundaries and TSSs not at boundaries. eRNA denotes the eRNA from non-genic boundaries, mRNA denotes the mRNA from genic boundaries **H.** Average predicted TTS sites from the TSSs identified in different classes of RNA by NET-CAGE. **I.** Average predicted 5’SS sites from the TSSs identified by NET-CAGE. The p-values in boxplots were calculated by the Wilcoxon rank-sum test. The boxplots depict the minimum (Q1-1.5*IQR), first quartile, median, third quartile and maximum (Q3+1.5*IQR) without outliers.

Next, we compared the stability, polyadenylation and splicing features of genic RNAs and eRNAs at TAD boundaries with those that are not within boundaries. In general, eRNAs exhibit fast turnover as seen by a higher ratio of nascent to mature eRNAs as compared to genic RNAs (37). By this measure eRNAs on non-genic boundaries were significantly more stable than other eRNAs, and RNAs on genic boundaries were slightly more stable than other genic RNAs (Fig. 3G).

The AATAAA polyadenylation motif is rare around Transcriptional Start sites (TSSs) of genic transcripts, increasing in abundance downstream as expected (Fig. 3H). Non-genic eRNAs on boundaries showed less enrichment of pre-mature AATAAA than other eRNAs (Fig. 3H), consistent with their higher stability. As expected, 5’ splice consensus sequences were strongly enriched in mRNAs relative to eRNAs (Fig. 3I), consistent with their greater stability. Together, enhanced stability, higher CTCF enrichment and insulation promoting activity of boundary eRNAs suggest that they have evolved to facilitate CTCF binding by direct physical interaction.

### Boundary RNAs enhance TAD transcription

The enhanced insulation we observed at sites of boundary eRNAs suggested that genes within the TADs of transcribed boundaries might be expressed at higher levels than those in TADs of non-transcribed boundaries. Accordingly, we compared the transcriptional levels of TADs with high, low, and randomly selected transcribed or non-transcribed boundaries. We found that the transcriptional output of entire TADs (Fig. 4A) or of individual genes within TADs (Fig. S7A) is correlated with the levels of RNAs at boundaries, and we obtained similar results for non-genic boundaries (Fig. S7B). To determine whether this positive association resulted from direct interactions between promoters and boundaries we analysed available promoter capture-seq data in HeLa cells (46). Boundaries were captured by promoters far more than by random regions (Fig. 4B), and genes that interacted with boundaries were relatively more active than genes that interacted with other genomic regions (Fig. 4C). Together, these data suggest that transcribed boundaries tend to interact with promoters and are associated with higher rates of TAD transcription. We confirmed the interaction of the *CDKN2A* promoter with the nearby transcribed boundary within the *INK4a/ARF* TAD using 4C-seq assays (Fig. 4D).

**Figure 4.**
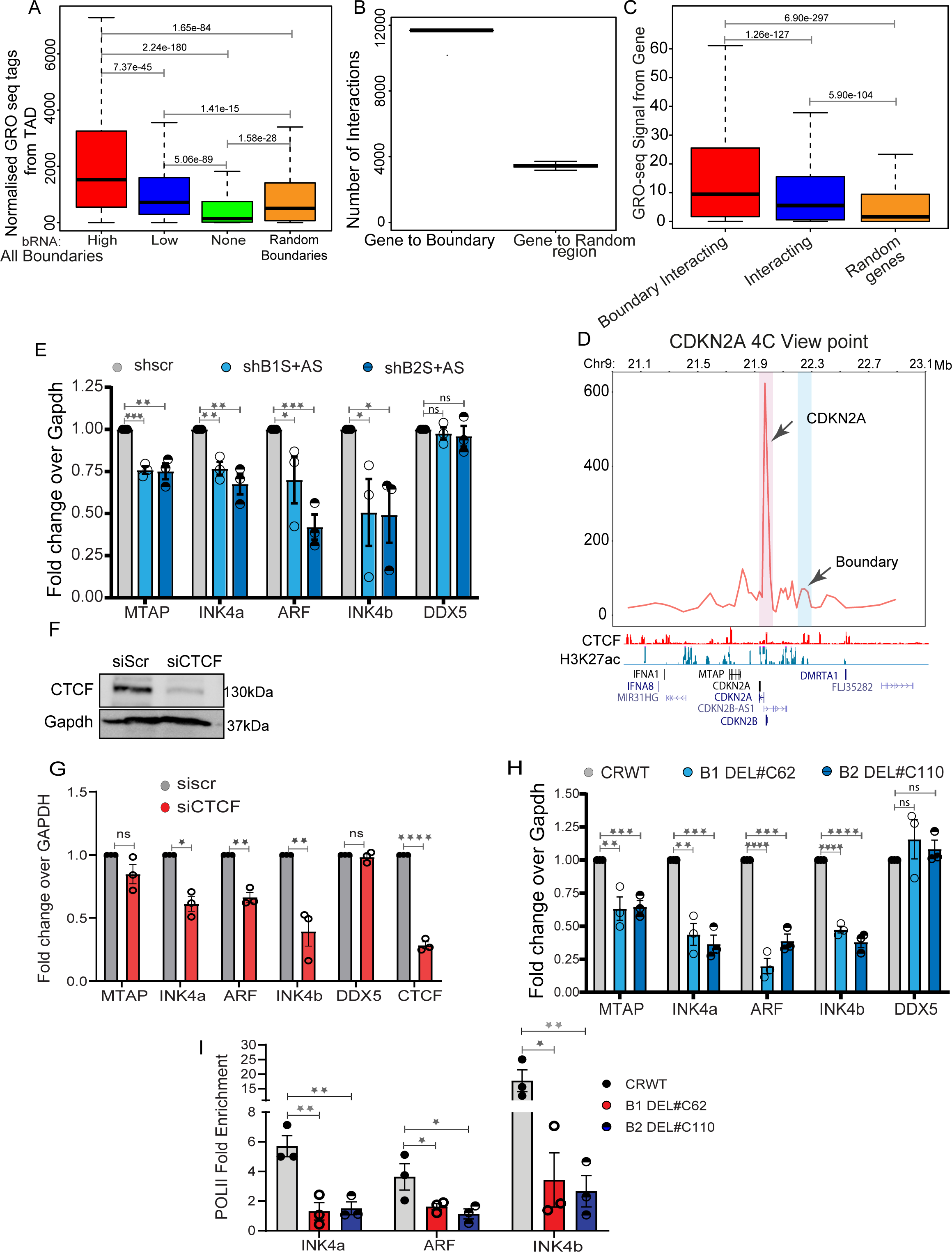
Boundary RNAs enhance TAD transcription: **A.** Box plots showing GRO-seq tags from the entire TAD with high, low, non-transcribed and random boundaries. **B.** Graph showing the number of interactions between genes and boundaries, and 100 randomizations of interactions between boundaries and random regions. **C.** Box plots showing the GRO-seq tag counts from genes when they interact with boundaries or any region vs. random interactions. **D.** 4C plot at the *CDKN2A* promoter viewpoint, where the interaction with the boundary is highlighted in blue. Below are H3K27ac, CTCF and gene annotations for HeLa cells. **E.** qRT-PCR plot shows the change in gene expression upon shRNA mediated knockdowns of boundary RNAs arising from B1 and B2 CTCF sites at the *INK4a/ARF* TAD boundary. **F.** Immunoblot showing the drop in CTCF protein levels upon its siRNA mediated knockdown. **G**. qRT-PCRs showing down-regulation of *INK4a/ARF* genes and levels of DDX5 and CTCF upon CTCF knockdown. **H.** qRT-PCR showing the changes in expression of different genes in the *INK4a/ARF* TAD and DDX5 upon Cas9-mediated independent deletion of B1 and B2 CTCF sites. **I.** PolII ChIP signal at *INK4a*, *ARF* and *INK4b* promoters in WT and in cells with B1 and B2 CTCF sites deletions. Error bars denote SEM from three biological replicates. *p*-values were calculated by Student’s two-tailed unpaired *t*-test in **E, G, H** and **I.** \*\*\*\**p* < 0.0001, \*\*\**p* < 0.001, \*\**p* < 0.01, \**p* < 0.05, ^ns^*p* > 0.05. p-values in boxplots were calculated by the Wilcoxon two-sided rank-sum test. The boxplots depict the minimum (Q1-1.5*IQR), first quartile, median, third quartile and maximum (Q3+1.5*IQR) without outliers.

Next, we asked if boundary RNAs play a direct role in transcriptional activity of genes within this TAD. Using shRNAs to sense and anti-sense B1 and B2 CTCF sites, we observed downregulation of *INK4a/ARF* (transcribed from *CDKN2A*), *INK4b* (transcribed from *CDKN2B*) and *MTAP*, whereas the transcription of *DDX5*, a non-*INK4a/ARF* TAD gene, remained unchanged (Fig. 4E). These data suggest that boundary eRNAs positively regulate the transcription of the genes within the *INK4a/ARF* TAD by direct physical interaction between promoters and boundaries.

Since the loss of specific boundary RNAs also caused loss of CTCF binding at the TAD boundary, we asked if the role of boundary RNA in TAD transcription is CTCF-dependent. Depletion of global CTCF using a pool of CTCF-specific siRNAs resulted in downregulation of genes within and outside of the *INK4a/ARF* TAD as expected (Fig. 4F-G). To test the effects of the loss of CTCF binding only at the *INK4a/ARF* TAD boundary, we deleted the B1 and B2 CTCF sites individually in HeLa cells (Fig. S8A-B) and observed downregulation of *INK4a/ARF* genes within the TAD (Fig. 4H) and the loss of PolII at promoters (Fig. 4I). We also observed that boundary eRNA knockdown and boundary deletion reduced the levels of the p16 protein encoded by *INK4a/ARF* (Fig. S8C). These data suggest that binding of CTCF mediated by boundary RNAs regulates the expression of genes within the *INK4a/ARF* TAD.

### Active enhancers within the TAD directly regulate boundary eRNA transcription

What is responsible for *INK4a/ARF* boundary transcription? Because HeLa cells are addicted to high levels of proteins encoded within the TAD (47-48), instead of deleting gene promoters, we used CRISPR inhibition (CRISPRi) using specific guide RNAs. We applied dCas9-KRAB on promoters of *CDKN2A*, *CDKN2B* and *MTAP* and assayed transcript levels at B1 and B2 CTCF sites (Fig. 5A). CRISPRi of these promoters resulted in transcriptional inhibition of these genes (Fig. 5B, red bars). However, boundary RNA levels were unaffected (Fig. 5B, blue bars). Consistent with unperturbed boundary RNA expression, there was no change in CTCF occupancy at B1 and B2 CTCF sites. However, we did observe loss of CTCF on the 5’ boundary that overlaps the MTAP gene, likely due to down-regulation of *MTAP* (Fig. 5C). The boundary of flanking TADs also did not exhibit any perturbations in CTCF occupancy (Fig. 5C). These data suggest that initiation of transcription from *CDKN2A* and *CDKN2B* promoters is not responsible for eRNAs at the boundary.

**Figure 5.**
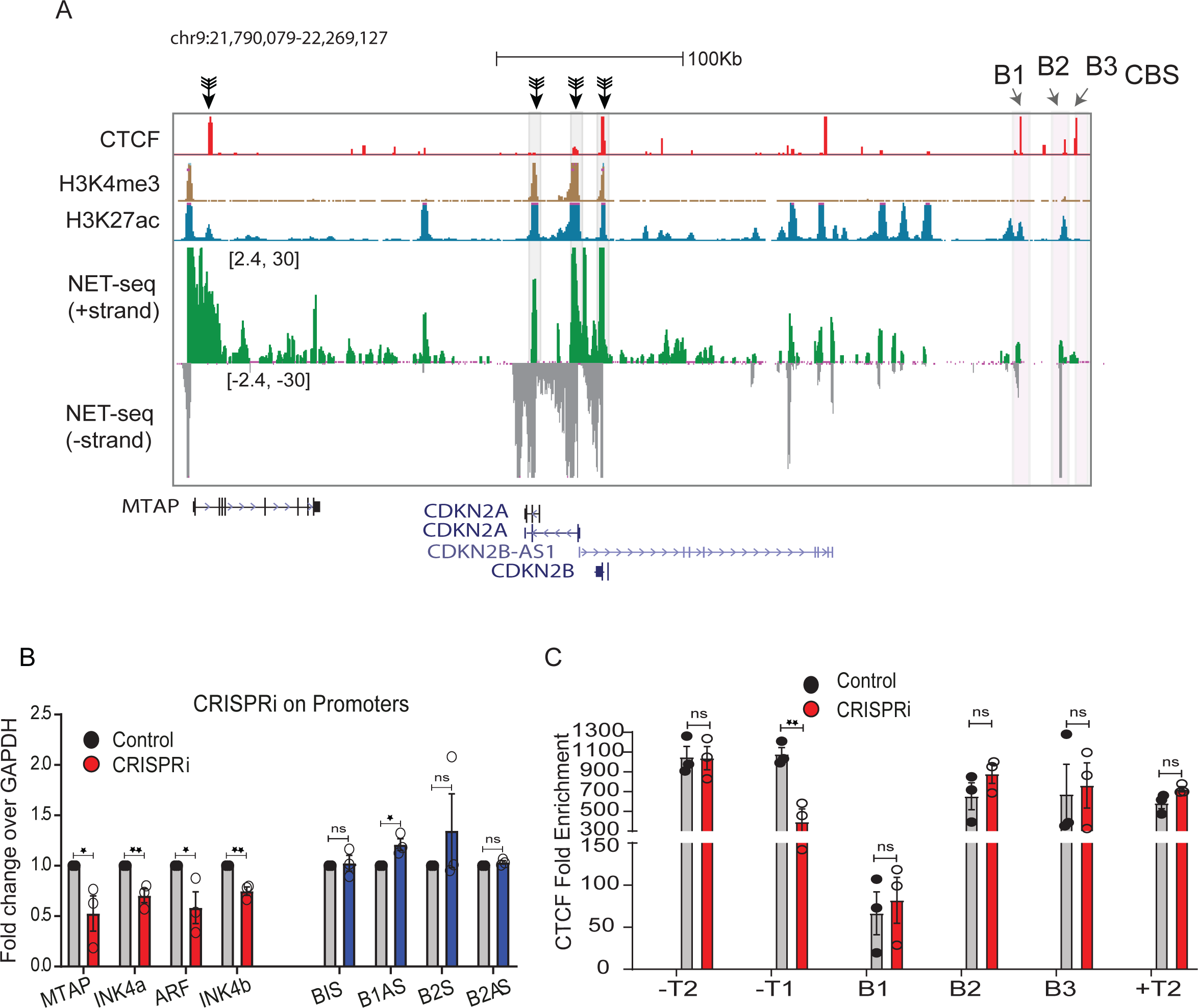
Transcriptional activity of promoters does not regulate non-genic transcription at boundaries: **A.** Browser shots displaying the promoters targeted by CRISPRi (Black arrows), the track is overlaid with CTCF, H3K4me3, H3K27ac and mNET-seq tracks. **B.** qRT-PCR assessing the changes in *INK4a/ARF* genes, sense and anti-sense boundary RNA arising from B1 and B2 CTCF sites upon CRISPRi on promoters. **C.** CTCF enrichment on boundaries of *INK4a/ARF* and its flanking TADs upon CRISPRi on *INK4a/ARF* promoters. Error bars denote SEM from three biological replicates. *p*-values were calculated by Student’s two-tailed unpaired *t*-test in **B** and **C.** \*\**p* < 0.01, \**p* < 0.05, ^ns^*p* > 0.05.

We next asked whether the *INK4a/ARF* TAD enhancers regulate genes within the TAD and boundary eRNAs. Genome-wide, transcribed boundary TADs are enriched in H3K27ac (Fig. 6A-B) and PolII (Fig. 6C) relative to non-transcribed boundaries and randomly selected genic or non-genic boundaries (Fig. 6C, Fig. S9A-B). We first tested the super-enhancer using E8, E12 and E17 enhancer deletion lines (Fig. 6D) in which *INK4a/ARF* genes were seen to be downregulated based on RNA-seq data (Fig. 6E) (41), confirming that these enhancers are functional enhancers. We then asked if these enhancers indeed regulate the boundaries as well. Interestingly, we observed down-regulation of sense and anti-sense boundary RNAs at the B1 and B2 CTCF sites upon E8, E12 or E17 deletions (Fig. 6F), which suggests that enhancers activate boundary transcription. To determine whether down-regulation might be caused by direct contact, we also performed 4C-seq on the E8 enhancer. We observed strong interactions with other enhancers in the super-enhancer cluster, with *CDKN2A* and *CDKN2B* promoters and with the boundary region (Fig. 6G, right highlighted pink). Importantly, deletion of the E8 enhancer conspicuously reduced 4C-seq signal, confirming that the interaction between the super-enhancer and distant B1 and B2 CTCF sites regulates boundary transcription (Fig. 6G). Although interactions between boundaries and enhancers have been previously reported (35) no regulatory relationship had been established, whereas our data demonstrate that apart from activating their target promoters, functional enhancers positively regulate the transcription at the boundaries by direct interactions.

**Figure 6.**
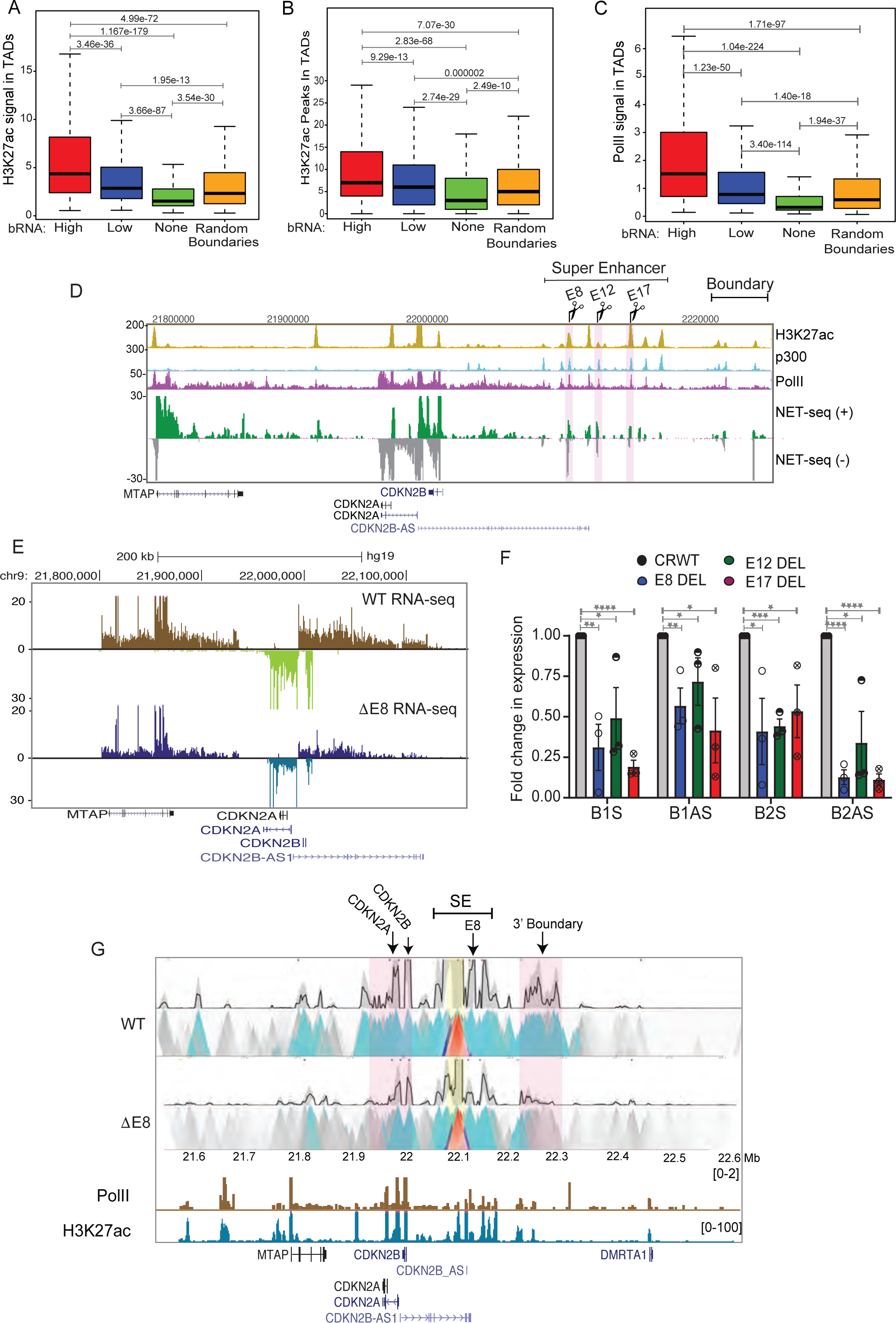
Active enhancers within a TAD directly regulate boundary eRNA transcription : **A.** Boxplots displaying enrichment of H3K27ac in TADs with high, low, non-transcribed and random boundaries. **B.** Boxplot showing number of H3K27ac peaks in TADs with various boundaries. **C.** Levels of PolII in TADs with different boundaries. **D.** UCSC browser shot showing H3K27ac, p300, PolII ChIP-seq and mNET-seq on the *INK4a/ARF* TAD. The tracks are overlaid by the gene annotations. The highlighted regions represent the enhancers (E8, E12 and E17) that were deleted. **E.** Browser shot shows RNA-seq at *INK4a/ARF* TAD in WT and E8 enhancer delete lines. **F.** qRT-PCR shows levels of sense and anti-sense boundary RNA arising from B1 and B2 CTCF sites upon E8, E12 and E17 enhancer deletions. **G.** 4C-seq plot in WT and in E8 deletion lines on an enhancer with the E8 enhancer as viewpoint (Yellow highlight) exhibiting interactions with promoters and 3’ boundary region (Pink highlight). Below are PolII, H3K27ac ChIP-seq tracks and gene annotations. Error bars denote SEM from three biological replicates. *p*-values were calculated by Student’s two-tailed unpaired *t*-test in **F.** \*\*\*\**p* < 0.0001, \*\*\**p* < 0.001, \*\**p* < 0.01, \**p* < 0.05. p-values in boxplots were calculated by the Wilcoxon rank-sum test. The boxplots depict the minimum (Q1-1.5*IQR), first quartile, median, third quartile and maximum (Q3+1.5*IQR) without outliers.

To ascertain whether high levels of boundary RNAs enhance insulation of the boundary *in vivo*, we interrogated the *INK4a/ARF* TAD upon senescence, where TAD transcription increases due to its cell cycle regulatory role. Upon senescence, bidirectional eRNA expression increases several-fold at the super-enhancer cluster (Fig. S10A, blue highlighted region), and *INK4a/ARF* gene expression and boundary RNA expression also increases (Fig. S10A, pink highlighted region). We validated these increases in gene transcription, boundary RNA expression and CTCF enrichment upon X-ray induced senescence (non-replicative senescence) (Fig. S10B-D), and also observed elevated p16 and p21 protein levels (Fig. S10E). Using HiC data, we observed increase in intra-TAD interactions at this TAD in senescent cells as compared to proliferative cells (Fig. S10F). Together these data suggest that increases in TAD enhancer activity accompany increases in their interactions with boundaries to induce boundary RNA expression, which in turn recruits more CTCF to enhance insulation.

### Boundary eRNAs strengthen TAD insulation

We noticed that TADs with transcribed boundaries exhibited higher strength of enhancer/promoter interactions than TADs with non-transcribed boundaries genome-wide (Fig. 7A). To test whether boundary function of the *INK4a/ARF* TAD affects super-enhancer/promoter interactions in a boundary RNA dependent manner, we performed 4C-seq assays at the E8 enhancer upon knockdown of sense and anti-sense boundary RNAs at the B1 CTCF site. As expected, we observed loss of boundary/enhancer interactions, but in addition, we observed weakening of *INK4a/ARF* enhancer/promoter interactions (Fig. 7B). Knockdown of B1 RNAs also resulted in increased interaction with the +T1 boundary of the neighboring TAD (Fig. 7B), which also showed a gain in CTCF binding (Fig. 2G). To confirm that boundary RNA loss was accompanied by reduced CTCF binding, we performed 4C-seq at the E8 enhancer using cells with a B1 CTCF site deletion. We again observed a loss of enhancer/promoter interaction and a concomitant increase in the interaction with the +1 T1 boundary of the neighboring TAD (Fig. 7B). These observations demonstrate that loss of a boundary RNA reduces CTCF binding and weakens insulation, resulting in aberrant enhancer interaction with an element outside of the TAD. We conclude that boundary RNAs are functional components of TAD boundaries (Fig. 7C).

**Figure 7.**
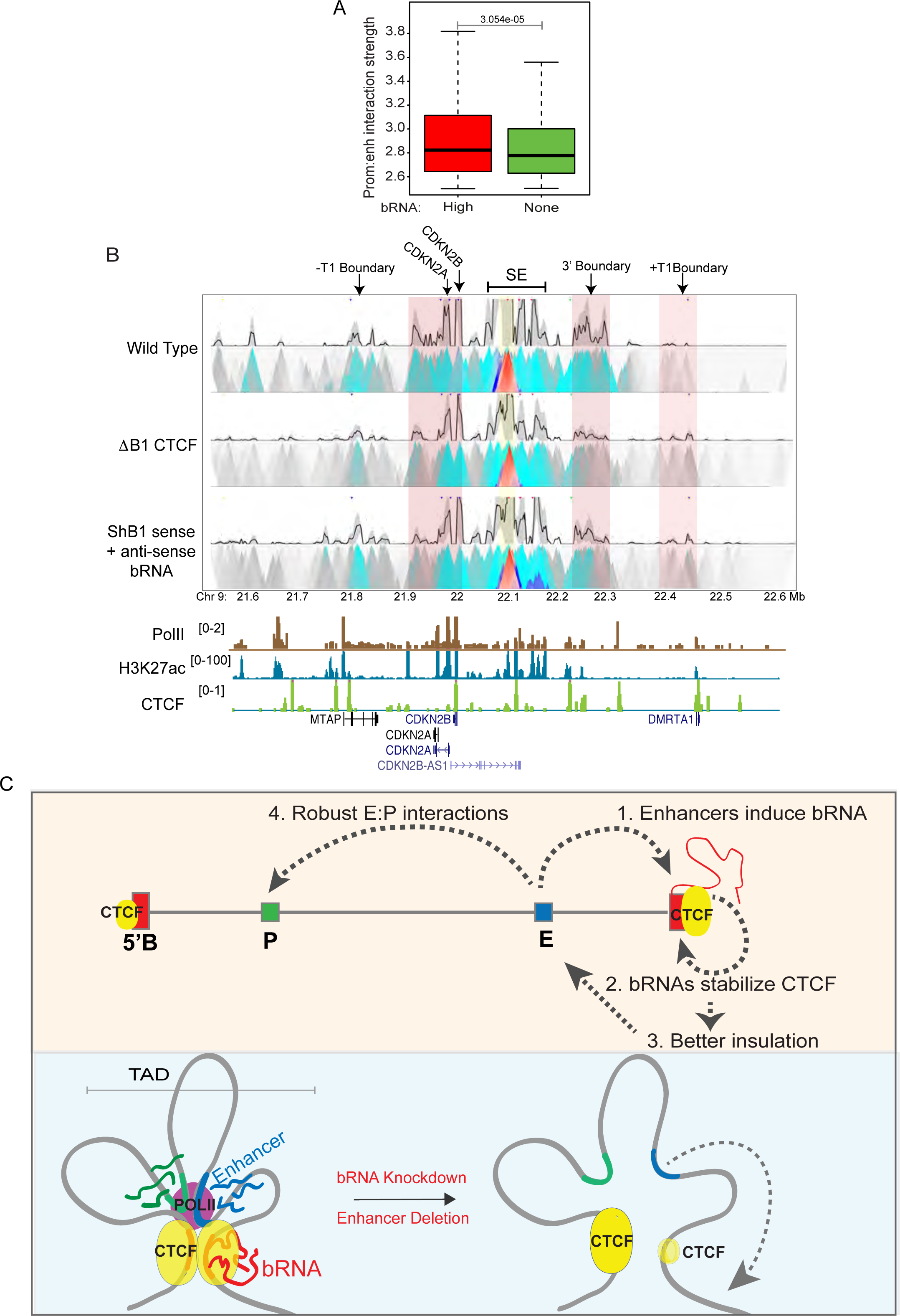
Boundary eRNAs strengthen TAD insulation: **A.** Enhancer:promoter interaction strength within TADs with high and non-transcribed boundaries. **B.** 4C-seq plot with the E8 enhancer as viewpoint (Yellow highlight) in WT and upon B1 CTCF site deletion (ΔB1) and knockdown of B1 sense and anti-sense boundary RNAs. The anchored region exhibit interactions with promoters and the 3’ and +T1 boundaries (Pink highlights). Below are PolII, H3K27ac and CTCF ChIP-seq tracks and gene annotations. **C.** Model: (1) An active enhancer physically interacts with a boundary and activates boundary RNA transcription. (2) The boundary RNA in return stabilizes the CTCF at these boundaries, thereby (3) strengthening insulation of these TADs. (4) This favors intra-TAD enhancer: promoter interactions to facilitate robust gene transcription. The loss of boundary RNA/enhancer deletion reduces the boundary RNA levels which triggers the loss of CTCF and insulation of TADs. Enhancer:promoter interactions weaken in these scenarios which causes concomitant loss of gene transcription. p-values in boxplots were calculated by the Wilcoxon rank-sum test. The boxplots depict the minimum (Q1-1.5*IQR), first quartile, median, third quartile and maximum (Q3+1.5*IQR) without outliers.

## Discussion

Interactions between CTCF and RNA have been implicated in genome organization (31-32), but the exact role of these interactions remains unknown. Here we show that eRNAs transcribed from TAD boundaries function in CTCF binding, TAD insulation and gene regulation. Our data provide direct evidence that boundary eRNAs create an insulated neighbourhood by associating with CTCF-bound sites and facilitating enhancer:promoter interactions for robust gene activation within a given TAD. We found that the transcriptional activity of promoters present within the TAD does not regulate non-genic boundary transcription. Using a series of enhancer and CTCF site deletions at the *INK4a/ARF* TAD, we have shown that enhancers interact with and positively regulate the expression of both genic and non-genic TAD boundary RNAs. Boundary RNAs, in turn, increase CTCF binding at TAD boundaries, strengthening weak CTCF motifs and increasing insulation. Transcription of boundary RNAs at the *INK4a/ARF* TAD is regulated by the super-enhancer within the TAD to facilitate CTCF binding and increased insulation at the boundary. This may explain why super-enhancers containing TADs exhibit stronger boundaries (34-36).

Boundary RNAs resemble eRNAs at active enhancers in being bidirectional (49-50), and as our study demonstrates, making direct contact with distant regulatory elements. However, unlike eRNAs, boundary RNAs contact enhancers that also contact promoters to enhance TAD gene activation. Popular chromosome conformation capture assays are strictly pairwise and cannot determine if enhancer elements activate promoters and boundary elements independently or together. However, more recently introduced 3D contact methods, including SPRITE (51) and GAM (52) are able to capture multiple simultaneous contacts, and in the future, may be applied to understand how enhancers regulate both gene promoter activity and transcription at boundaries to strengthen TADs.

Furthermore, in the absence of the RNA binding domain of CTCF, cells exhibit a decrease in the total number of TADs and an increase in the size of individual TADs due to reduced insulation between neighboring TADs (31). A clear mechanistic basis for this observation comes from our findings in which TAD enhancer-mediated CTCF-RNA interactions maintain strong TAD boundaries by increasing insulation between neighboring TADs. Thus, the inability of CTCF mutants to interact with RNA will compromise CTCF enrichment and its insulating function at TAD boundaries, leading to the merging of adjacent TADs into larger TADs. In such cases, less CTCF at the boundary might allow the cohesin ring to slip to the next TAD thereby compromising the insulation (53-54). Thus a positive feedback loop is established whereby super-enhancers maintain high specificity for promoters within the TAD by also activating boundary transcription and preventing interactions with neighboring promoters.

TAD boundaries are enriched for H3K27ac and PolII, suggesting the presence of active regulatory elements and transcription (24,34-36). However, activation of transcription at the boundary alone is not sufficient to maintain a TAD boundary (23,55). Our study shows that the products of the TAD boundary transcription *i.e.*, boundary RNAs, are required to strengthen TAD insulation by increasing CTCF occupancy. Transcription-assisted TAD boundary strengthening may also be a factor in zygotic activation-dependent boundary formation where early transcribing genes serve as the nucleation sites for TAD boundaries (55). Similarly, transcription and the resulting RNA may be involved in forming distinct TADs boundaries around transcribed escape genes on the inactive chromosome during X chromosome inactivation (26).

## Materials and Methods

### Cell Culture

HeLa cells were obtained from ATCC and were cultured in (DMEM) Dulbecco’s Modified Eagle’s medium supplemented with 10% Fetal Bovine Serum and 1% Penicillin/Streptomycin. The cells were maintained at 37°C in the presence of 5% CO2 in a humidified incubator.

### Chromatin Immunoprecipitation

Cells were crosslinked with 1% formaldehyde at room temperature for 10 min with constant shaking. Glycine was added to a final concentration of 125 mM to quench the formaldehyde. Cells were washed three times with 1X ice-cold PBS then were scraped and pelleted down 1X PBS at 4 °C. Cells were gently resuspended in nuclear lysis buffer (50 mM Tris-HCl pH 7.4, 1% SDS, 10 mM EDTA pH 8.0) supplemented with 1X PIC and incubated on ice for 10 min. The lysate was subjected to fragmentation using a Diagenode Bioruptor Pico for 20 cycles (30 s ON and 30 s OFF) to generate fragments of around ∼500 bp. The cell lysate was cleared at 12K rpm for 12 min. For each IP, 100 µg of sheared chromatin was used. The lysates were diluted by adding Dilution Buffer (DB) (20 mM Tris-HCl pH 7.4, 100 mM NaCl, 2 mM EDTA pH 8.0, 0.5% Triton X-100) supplemented with 1X PIC in 1:1.5 ratio (1 volume of sheared chromatin and 1.5 volumes of Dilution Buffer). One µg of antibody was added to immunoprecipitate the DNA and incubated on a rocking platform overnight at 4°C. Fifteen µl of Pre-blocked (with 1% BSA) Protein G Dynabeads (10004D, Invitrogen) was added to the tubes and rotated for 4h at 4 °C. Beads were collected, flow-through was discarded and the samples were washed with Wash Buffer I (20 mM Tris-HCl pH 7.4, 150 mM NaCl, 0.1% SDS, 2 mM EDTA pH 8.0, 1% Triton X-100) at 4 °C on a rocking platform. Washes were sequentially repeated with Wash Buffer II (20 mM Tris-HCl pH 7.4, 500 mM NaCl, 2 mM EDTA pH 8.0, 1% Triton X-100), Wash Buffer III (10 mM Tris-HCl pH 7.4, 250 mM LiCl, 1% NP-40, 1% Sodium Deoxycholate, 1 mM EDTA pH 8.0 and 1X TE (10 mM Tris pH 8.0, 1mM EDTA pH 8.0). Chromatin was eluted with 200 ul of Elution Buffer (100 mM NaHCO3, 1% SDS) for 30 min at 37°C in a thermomixer. The supernatant was collected in separate tubes and 14 µl NaCl (5 M) was added to eluted samples and kept at 65 °C overnight for de-crosslinking. Immunoprecipitated DNA was purified by phenol:chloroform:isoamyl alcohol, followed by ethanol precipitation. The final air-dried DNA pellet was dissolved in 100 µl of 1X TE. These samples were then used for qRT-PCRs using SYBR Green. Mean values for all the regions analyzed in different conditions were expressed as fold-enrichment compared over beads. Statistically significant differences between different conditions were computed using a *T*-test (*p*-value *<* 0.05), ChIP primers are listed in Supplementary Table. 1.

### RNA Isolation and cDNA synthesis

Cells were lysed in 1 ml of Trizol (Thermo Fisher Inc.). To each sample, 200 µl chloroform was added, briefly vortexed and centrifuged at 12K rpm for 12 min. The aqueous phase was carefully collected and transferred to the fresh tube. One volume of isopropanol was added to the sample and incubated at room temperature for 10 min to precipitate the RNA. The samples were centrifuged at 12K rpm for 12 min, and supernatants were discarded without disturbing the pellet. The pellet was washed with 75% ethanol. The pellet was air-dried and dissolved in RNase-free water. The RNA was treated with ezDNase (Invitrogen) to remove the traces of contaminating DNA. One µg of RNA was used for each cDNA synthesis reaction by Superscript IV (Invitrogen) and random hexamers as per the manufacturer’s recommendation. The CFX96 Touch (Biorad) real-time PCR instrument was used for qRT-PCRs. qRT-PCRs were performed using three biological replicates for each sample. Fold change was calculated by the ΔΔCt method and individual expression data were normalized to GAPDH mRNA. The *p*-values were calculated by Student’s unpaired two-tailed *t*-test. qRT-PCR primers are listed in Supplementary Table 1.

### CRISPR Cas9 mediated deletion

Guide RNAs (gRNAs) were designed with the crispr.mit.edu tool. gRNAs were selected based on the highest score and the least number of off-targets. gRNAs were cloned in a pgRNA humanized vector (Addgene #44248, a gift from the Stanley Qi Lab) between BstX1 and Xho1 restriction sites. gRNAs were co-transduced with a lenti-Cas9 vector. Cells were placed under selection using puromycin (3µg/ml) for 48 hours. Single cells were seeded in a 96 well plate. Wells with single cells were marked and allowed to grow until they formed colonies. The cells were trypsinized and shifted to 48 well plates and allowed to grow to confluence, and then half of the cells were taken for the Surveyor assay and the other half was plated again. Surveyor assays were performed using a PCR-based method using primers listed in the Supplementary Table 1.

### CRISPRi gRNA design and cloning

We used GPP sgRNA Designer (https://portals.broadinstitute.org/gpp/public/analysis-tools/sgrna-design) to design the gRNAs against TSSs of *MTAP*, *CDKN2A* and *CDKN2B* in the *INK4a/ARF* TAD. For each gene, two gRNAs were designed. The gRNAs were cloned in a customized PX459 vector (pSpCas9(BB)-2A-Puro V2.0, Addgene #62988, a gift from the Feng Zhang lab). The Cas9 enzyme cassette was removed from PX459 by digestion with Xba1 and Not1 followed by blunt-end ligation. The cloning of the gRNAs in the customized PX459 plasmid was performed as per the Zhang Lab’s general cloning protocol. The dCas9 plasmid, together with gRNA specific to targeted genes, were transfected in 70% confluent cells. Transfections were done using Lipofectamine 2000 (Invitrogen, cat no #11668027). CRISPRi cells were harvested for ChIP and gene expression analysis.

### shRNA design and transfection

We used an shRNA design tool (https://portals.broadinstitute.org/gpp/public/seq/search) to design the shRNAs against RNA at B1 and B2 CTCF sites within the *INK4a/ARF* TAD boundary. The shRNAs were cloned in a customized pLKO.1 puro, (Addgene #8453, a gift from Bob Weinberg. The cloning of the shRNAs in the pLKO.1 plasmid was performed as per the Addgene’s pLKO.1 cloning protocol. shRNA-specific plasmid together with lentiviral packaging plasmids like VSVG (a gift from Bob Weinberg, Addgene #8454), and PAX2, a gift from Didier Trono (addgene #12260) were co-transfected using Lipofectamine 2000 Invitrogen (cat #11668027). Cells were selected by puromycin (3µg/ml) (GIBCO cat # A11138-03) and knockdowns were confirmed using qPCR oligos listed in Supplementary Table 1.

### siRNA transfection

SMARTpools was used to design scrambled (cat no D-001810-10-05) and CTCF siRNAs (L-020165-00-0005) were purchased from GE Dharmacon. Transfections were done using Lipofectamine 2000 (Invitrogen cat no 11668027).

### Lentiviral transduction

HEK293FT cells were grown in poly-D lysine coated culture dishes. These cells were co-transfected with lentiviral packaging plasmids like VSVG (a gift from Bob Weinberg, Addgene #8454) and PAX2 (a gift from Didier Trono, Addgene #12260) along with the plasmid of interest using Lipofectamine 2000 Invitrogen (cat# 11668027). The medium was changed after 6 hours. The viral soup was collected after 48 h and 72 hours, pooled together, filtered with a 0.44 µm syringe filter and then finally added to cells along with 8 µg/ml of polybrene. Transduction was stopped after 16 hours.

### RNA Immunoprecipitation

For RNA, primers were designed to amplify desired genomic regions that correspond to peaks of boundary RNA (GRO-seq) at B1 and B2 CTCF sites of the *INK4a/ARF* TAD boundary. The amplified products were cloned into the pcDNA3 BoxB plasmid (a gift from Howard Chang, Addgene #29729). All clones were confirmed by Sanger sequencing and subsequently used for RNA synthesis using T7 RNA polymerase (Promega Inc. P207e) and biotin RNA labeling mix (Roche 11685597910). Primer sequences are listed in Supplementary Table 1.

RNA immunoprecipitation was adopted from (56), with some modifications. HeLa cells were grown to 90–95% confluency and were harvested by scraping and washed twice with ice-cold 1X PBS to gently resuspend in 2 ml of PBS. A volume of 2 ml of nuclear isolation buffer [NIB: 40 mM Tris–HCl pH 7.5, 20 mM MgCl_2_, 1.28 M sucrose, 4 % Triton X-100, 1 mM PMSF, protease inhibitors, and 20 U/ml SUPERase inhibitor (Thermo Fisher Scientific, catalog # AM2694)] was added and the pellet was gently resuspended. Then, 6 ml of distilled water was added and kept on ice for 20 min with intermittent gentle shaking. Nuclei were pelleted by centrifugation at 2500×*g* for 5 min at 4°C. The pellet was resuspended in 1 ml of RIP buffer (25 mM Tri–HCl pH 7.4, 150 mM KCl, 0.5 mM DTT, 0.5 % NP40, 1 mM PMSF, protease inhibitors, and 20 U/ml SUPERase inhibitor) and incubated on ice for 5 min. Nuclei were sheared by 10 cycles of sonication (30s on and 30 s off) in a Bioruptor (Diagenode) followed by centrifugation at 12K rpm for 10 min at 4°C. One µg of biotinylated RNA was incubated with 20 µl of RNA structure buffer (RSB) (10 mM Tris–HCl pH 7.0, 100 mM KCl, 10 mM MgCl_2_, 1 mM PMSF, protease inhibitors, and 20 U/ml SUPERase inhibitor) at room temperature for 5 min. Folded RNA was mixed with 1 mg of nuclear extract in 500 µl of RIP buffer and rotated at 4^0^C for 1h. Fifteen µl of Dynabeads MyOne Streptavidin beads was added and rotation continued for one more hour. Samples were washed three times with RIP buffer and beads with proteins were boiled in 2XSDS for 10 min. Immunoblotting for CTCF was performed using a CTCF antibody (Cell Signaling Technology, #3418).

### UV RIP

Ultraviolet-RNA Immunoprecipitation was performed as in (57) with some modifications. HeLa cells at a confluency of 80–90% were cross-linked by UV irradiation in Stratalinker UV cross-linker. Cells were washed three times with cold 1XPBS and scraped in PBS before subjecting to centrifugation at 2500 rpm. Pellets were lysed in RIP lysis buffer [25 mM HEPES-KOH at pH 7.5, 150 mM KCl, 0.5% NP40, 1.5 mM MgCl_2_, 10% glycerol, 1 mM EDTA, 0.4 U RNase inhibitor (Invitrogen, 18091050)], and 1X protease inhibitor cocktail (PIC) on ice for 30 min. Cleared cell lysates were subjected to IP with CTCF antibody-bound Protein G Dynabeads (Invitrogen, 10004D) overnight. Samples were subsequently washed three times with RIP lysis buffer and RNA samples were eluted using TRIzol reagent. Prior to cDNA synthesis, RNA was treated with RNase-Free DNase Qiagen (cat no.79256) to remove the traces of contaminating DNA. cDNA was prepared using random hexamers by Superscript IV (Invitrogen 18091050) as per the manufacturer’s recommendation and analyzed by PCR primers listed in Supplementary Table 1 and were run on 1% Agarose gel.

### Cell Fractionation

Cell fractionation and RNAseA treatment of nuclei was adapted from (58). Briefly, cells were washed three times with cold 1XPBS and scraped in 1ml of PBS, pelleted down at 2500 rpm for 5 min at 4^0^C. Pellets were then resuspended in Hypotonic Lysis Buffer (HLB) (10 mM Tris HCl pH 7.5, 10 mM NaCl, 3 mM MgCl_2_, 0.3% NP-40, 10% glycerol, 1XPIC), the mix was incubated on ice for 10 min and then centrifuged at 800g for 8 min at 4^0^C. The supernatant was transferred to a new tube and marked as Cytoplasmic fraction. The nuclei pellet after centrifugation was washed twice with HLB and centrifuged at 200g for 2 min at 4^0^C. The pellet was then resuspended in 700 ul of ice cold Modified Wuarin-Schiebler buffer (MWS: 10 mM Tris-HCl, pH 7.5, 300 mM NaCl, 4 mM EDTA, 1 M urea, 1% NP-40, 1% glycerol, 1XPIC) with or without 0.1l of 100mg/ml RNaseA (Qiagen cat. no. 19101) for 15 min on ice briefly tapped few times during incubation time and then centrifuged at 1000g for 5 min at 4^0^C. The supernatant was collected as a nucleoplasmic fraction. The chromatin pellet was washed twice with MWS buffer and centrifuged at 500g for 3min at 4^0^C. The pellet was then dissolved in 500 µl of Nuclear Lysis Buffer (NLB: 20 mM Tris HCl pH 7.5, 150 mM KCl, 3 mM MgCl_2_, 0.3% NP-40, 10% glycerol, 1XPIC) and sonicated for 10 cycles (30 s on, 30 s off) in a Bioruptor (Diagenode). SDS loading dye was added and the samples were boiled before immunoblotting.

### Chromatin-associated RNA isolation

Chromatin-associated RNA isolation was adapted from (59). Trypsinized cells were collected in 1.5 ml tubes and washed twice with 1xPBS and pellet down at 200xg for 2 min. To each pellet was added 400 µl of Cell lysis buffer (10 mM Tris HCl ph7.4, 150 mM NaCl, 0.15% Igepal), gently pipetted 3-5 times and then incubated for 5 min on ice. To the Lo-bind 1.5ml tube was added 1 ml (2.5 volume) of cold sucrose buffer (10mM Tris HCl ph7.4, 150mM NaCl, 24% Sucrose) and gently overlaid with the cell lysate followed by centrifugation at 3500xg for 5 min to collect the nuclear pellet. Isolated nuclei were rinsed with 1 ml of ice-cold 1xPBS-EDTA solution followed by a short spin at 3500xg. The nuclei were resuspended in 250 µl of glycerol buffer (20 mM Tris HCl ph7.4, 75 mM NaCl, 0.5 mM EDTA pH 8.0, 50% Glycerol) and then immediately 250 µl of Urea buffer (10 mM Tris HCl ph7.4, 1 M Urea, 0.3 M NaCl, 7.5 mM MgCl_2_, 0.2 mM EDTA, 1% Igepal) was added, mixed by vortexing for 4 s and then incubated on ice for 2 min. Chromatin was centrifuged at 13000xg for 2 min to collect the Chromatin-RNA complex. The chromatin-RNA pellet was washed with 1xPBS-EDTA once and RNA was isolated using Trizol Reagent. RNA obtained was treated with ezDNase (Ambion) to remove the traces of contaminating DNA. One µg of RNA was used for each cDNA synthesis reaction by Superscript IV (Invitrogen) and random hexamers as per the manufacturer’s recommendation. The CFX96 Touch (Biorad) real-time PCR was used for qRT-PCRs. qRT-PCRs were performed using three biological replicates for each sample. Fold changes were calculated by the ΔΔCt method and individual expression data was normalized to pre-c-myc. The *p*-values were calculated by Student’s unpaired two-tailed *t*-test. qRT-PCR primers are listed in Supplementary Table 1.

### X-ray induced Senescence

BJ fibroblasts were cultured in (DMEM) Dulbecco’s Modified Eagle’s medium supplemented with 10% Fetal Bovine Serum and 1% Penicillin/Streptomycin at 37 °C in a humidified incubator containing 5% CO2. Cells were first immortalized by expressing hTERT. Cells were then seeded at around 40% confluency. The day after seeding, a single dose of 10 Gy was delivered using a CIX3 cabinet x-ray irradiator (xstrahl Life Sciences) (195 kV, 10 mA). Irradiated cells were not passaged, but the medium was changed on Days 4, 8,12 and 16 and cells were either harvested on day 10 or day 20 and used for RNA isolation and Chip experiments respectively. Senescence was confirmed by SA-β-gal staining (Cell Signaling technology #9860) and p21 expression (p21 Santacruz Biotechnology sc-397).

### Antibodies

Antibodies used were: CTCF (Cell Signaling Technologies, #3418 for immunoblotting and Millipore Sigma #07-729 for CUT&RUN.RNase), DDX5 (Abcam, ab-21696), Cdkn2A, p16 (Abcam ab-108349), ARF, p14 (Santacruz Biotechnology, sc-53639), PolII (Santacruz Biotechnology, sc-55492), GAPDH (Santacruz Biotechnology, sc-32233), Histone H3 (Sigma-Aldrich H0164).

### ChIP-seq and CUT&RUN analysis

Sequenced reads were aligned to the hg19 assembly using default Bowtie2 options. Tag directories were made from the aligned reads using the HOMER (http://homer.ucsd.edu/homer/) makeTagDirectory command. A 200 bp sliding window was used to identify narrow peaks of transcription factors’ ChIP-seq and CTCF CUT&RUN. CUT&RUN.RNAse was performed as described in (26). Artifacts from clonal amplification were neglected as only one tag from each unique genomic position was considered (-tbp 1). The threshold was set at a false discovery rate (FDR) of 0.001 determined by peak-finding using randomized tag positions in a genome with an effective size of 2 × 10^9^ bp. For histone marks, seed regions were initially found using a peak size of 500 bp (FDR<0.001) to identify enriched loci. Enriched regions separated by <1kb were merged and considered as blocks of variable lengths. The HOMER makeUCSCfile command was used to generate bedgraph files of the read densities across the genome and this track was uploaded to the UCSC genome browser. The tag directories generated were used to quantify signals at various regions of interest using annotatePeaks.pl. The datasets used in the study are listed in Supplementary Table 3.

### GRO-seq analysis

The sequenced reads were aligned to the hg19 assembly using default Bowtie2 options. Tag directories were made from the aligned reads using the HOMER makeTagDirectory command. Tag directories were used to quantify signal and gene expression. The HOMER makeUCSCfile command was used to generate strand-specific bedgraph files of the read densities across the genome and this track was uploaded to the UCSC genome browser.

### TADs and loop domain calling

Raw paired-end reads were trimmed and individually mapped to the hg19 assembly using bowtie2. The HOMER makeTagDirectory was first used to create tag directories with tbp 1 option. Datasets were further processed by HOMER in order to remove small fragment and self-ligations using makeTagDirectory with the following options: -removePEbg -removeSpikes 10000 5. Next, findTADsAndLoops.pl was used to obtain overlapping TADs (TADs and sub-TADs), produced at 5 kb resolution with 10 kb windows. This program was also used to generate two bedGraph files that describe the directionality index and the insulation score. tagDir2hicFile.pl was used to generate .hic files. The WashU epigenome browser was used to visualize the HiC data. Boundaries of the TADs were defined as 5 kb or 20 kb regions centered at the end of the TADs called. HICCUPS was used to obtain loop anchors from the .hic file at a 5kb resolution. HOMER annotatePeaks.pl was used to quantify signal from tagDirectories, count peaks, quantify insulation and annotate features at the TAD boundaries. Non-genic TAD boundaries were identified as those that did NOT intersect with an annotated gene body. GRO-seq signal was used to rank the boundaries and to then select 10% of unique boundaries in each category (high transcription, low transcription, no transcription and random). The .hic file was converted to .cool. The file was then used to plot the averaged insulation score valleys at 5 kb resolution and 100 kb around the boundary using coolpup.py.

### NET-CAGE and CAGE Analysis

NET-CAGE and CAGE sequenced reads were aligned to hg19 using bowtie2. The HOMER program makeTagDirectory was first used to create tag directories. findPeaks with “–style tss” was used to obtain the NET-CAGE TSSs. TSSs identified within 1 kb of each other on the same strand were merged. NET-CAGE identified TSSs intersecting the refseq annotated TSSs were called mRNA TSSs and the rest were defined as eRNA TSSs. Those TSSs which fell in boundaries were identified as bmRNAs or beRNAs. A span of 2kb around each identified TSSs was used to quantify NET-CAGE and CAGE signal and to calculate the log2 fold difference between the two.

### 5’ Splice site and Transcription Termination Site Sequence Analysis

To identify the 5’ SS motif, HOMER denovo motif analysis was performed on the 500bp region after the Refseq annotated TSSs with other H3K27ac 500 bp regions as background. The top hit was observed to be the splice acceptor site. The “AATAAA” motif was used as the 3’ polyA sequence. HOMER scanMotifGenomeWide was used to identify “predicted” splice sites and polyA terminator sites using previously identified motifs. The occurrence of predicted sites was plotted at the previously identified TSSs.

### Circular Chromatin Conformation Capture-seq

4C was performed as per the protocol described in van de Werken et al., 2012 with minor modifications. HeLa cells were fixed with fresh formaldehyde (1.5%) and quenched with glycine (125mM) followed by washes with ice-cold PBS (2x) and scraped, pelleted and stored at -80^0^C. Lysis buffer [Tris-Cl pH 8.0 (10mM), NaCl (10mM), NP-40 (0.2%), PIC (1x)] was added to the pellets and homogenized with a Dounce homogenizer (20 stroked with pestle A followed by pestle B). The 3C digestion was performed with HindIII (400units, NEB) and ligation was performed using the T4 DNA ligase and ligation mix (1% Triton X-100, 0.105 mg/ml BSA, 1.05 mM ATP, 1x Ligation buffer, where 10x Ligation buffer is 500 mM Tris-Cl pH 7.5, 100 mM MgCl_2_, 100 mM DTT). The ligated samples were then purified by PCI (Phenol/Chloroform/Isoamyl alcohol) extraction, subjected to ethanol precipitation, and the pellet was eluted in TE (pH 8.0) to obtain the 3C library. The 4C digestion was performed using 50 units DpnII (NEB) and the samples were ligated, purified and precipitated similar to the 3C library to obtain the 4C library. The 4C library was subjected to RNaseA treatment and purified by the QIAquick PCR purification kit. The concentration of the library was then measured by Nanodrop and subjected to PCR using the oligos for the enhancer viewpoint. The samples were next PCR purified using the same kit and subjected to Illumina HiSeq2500 sequencing using 50 bp single-end reads.

### 4C method and analysis

The sequenced reads were aligned to the hg19 assembly as described in R.4Cker (https://github.com/rr1859/R.4Cker) with Bowtie2. Data analysis was further performed using 4C-ker with default parameters. 4Cseqpipe (http://compgenomics.weizmann.ac.il/tanay/?page_id=367) was also used to process the sequenced data. 4C-seq images were generated using truncated mean at a 10kb resolution.

### Motif analysis

annotatePeaks.pl was used with –hist option with the CTCF motif probability matrix to plot the motif occurrence in the peaks.

### Promoter capture analysis

Processed CHiCAGO data was obtained from Thiecke et al. (https://osf.io/brzuc/). The BEDTools suite was used to obtain interactions of interest and perform further analysis. The promoter interacting region of the Bedpe files of pcHiC interactions were then intersected with boundaries to obtain promoters that interact with boundaries. The strength of the interactions obtained from the processed CHiCAGO data was plotted for interactions from boundary interacting promoters to enhancers (H3K27ac regions).

### RNA-seq analysis

The raw reads were mapped to the hg19 assembly using hisat2 in a strand-specific manner. Tag directories were made from the aligned reads using the HOMER makeTagDirectory command.

The HOMER makeUCSCfile command was used to generate strand-specific bedgraph files of the read densities across the genome and this track was uploaded to the UCSC genome browser.

## Acknowledgments

We are grateful to Henrietta Lacks and to her family members for their contributions to biomedical research. We thank DN lab members, Ranveer Jayani, Amanjot Singh for the discussions.

## Author contributions

Conceptualization and design: ZI, BS and DN NGS analysis: BS

Methodology: ZI, BS, UF and KW CTCF CUT&RUN: JT and SH NGS library preparation and sequencing: AP Writing—original draft: DN, SH, ZI and BS

Reviewing and editing: DN, ZI, BS, KW, UF, JT, AKS, AP, RS

## Data and materials availability

The accession codes for data used in the study are mentioned in Table. 3. The CUT&RUN raw and processed data for control and RNaseA-treated HeLa cells have been deposited in the GEO repository with accession code GSE180659. The data can be reviewed using token ehwnowswbbcdxyl into the box.

Goto https://www.ncbi.nlm.nih.gov/geo/query/acc.cgi?acc=GSE180659

---

**Supp Figure 1.**
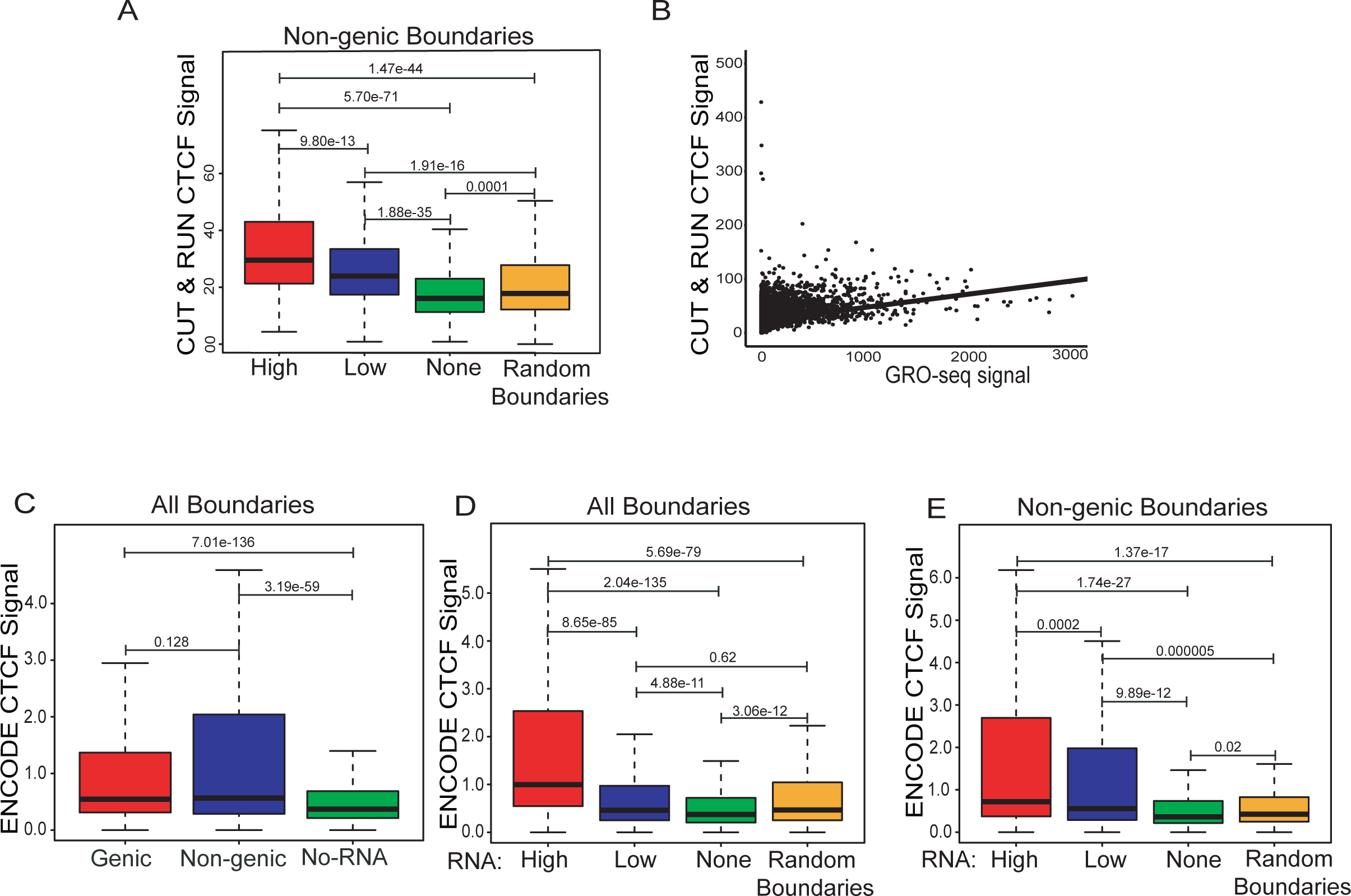
Transcribed boundaries exhibit better insulation: **A.** CTCF enrichment (CTCF CUT&RUN) at non-genic transcribed boundaries with high, low RNA versus non-transcribed and random boundaries **B.** levels of RNA and CTCF occupancy at TAD boundaries are positively correlated. **C.** CTCF enrichment (ChIP-seq) at transcribed genic, non-genic and non-transcribed boundaries. **D.** CTCF enrichment at all transcribed boundaries with high, low RNA versus non-transcribed and random boundaries. **E.** CTCF enrichment at non-genic transcribed boundaries with high, low non-coding RNA versus non-transcribed and random boundaries. p-values in boxplots were calculated by the Wilcoxon rank-sum test. The boxplots depict the minimum (Q1-1.5*IQR), first quartile, median, third quartile and maximum (Q3+1.5*IQR) without outliers.

**Supp Figure 2.**
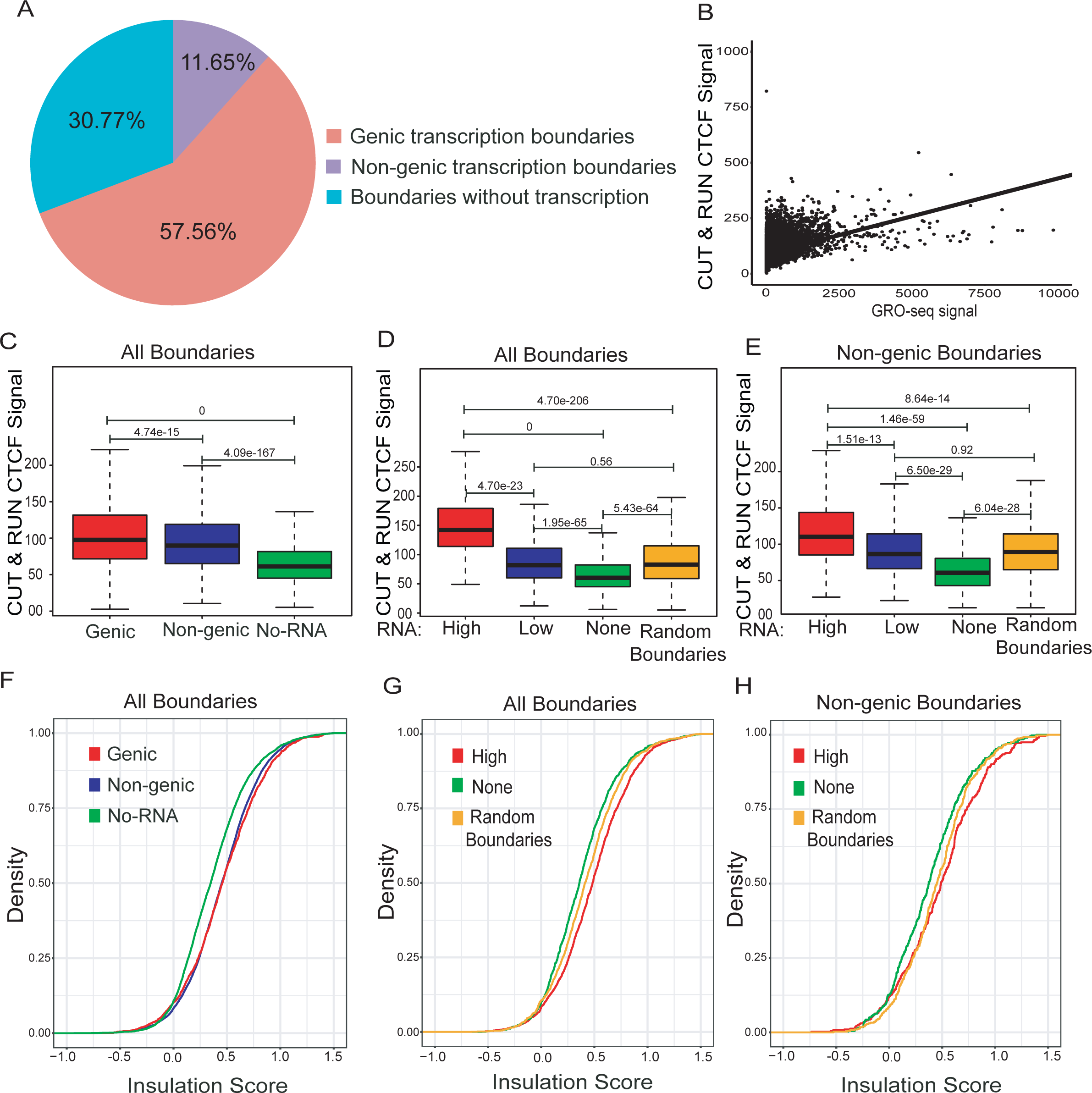
boundaries defined at 20 kb resolution show the similar features as at 5 kb resolution: **A.** Pie chart displaying the percentage of boundaries that show genic and non-genic transcribed boundaries and not transcribed boundaries. **B.** Levels of RNA and CTCF occupancy at TAD boundaries are positively correlated. **C.** CTCF enrichment on transcribed genic, non-genic and non-transcribed boundaries. **D.** CTCF enrichment on all transcribed boundaries with high and low RNA versus non-transcribed and random boundaries. **E.** CTCF enrichment on non-genic transcribed boundaries with high and low non-coding RNA versus non-transcribed and random boundaries. **F.** Density plot showing insulation scores for transcribed genic, non-genic and non-transcribed boundaries. **G.** Density plot showing insulation scores for all transcribed boundaries exhibiting varying levels of RNA vs. non-transcribed and random boundaries. **H.** Density plot showing insulation scores of non-genic transcribed boundaries exhibiting varying levels of RNA versus non-transcribed and random boundaries. p-values in boxplots were calculated by Wilcoxon rank-sum test. The boxplots depict the minimum (Q1-1.5*IQR), first quartile, median, third quartile and maximum (Q3+1.5*IQR) without outliers.

**Supp Figure 3.**
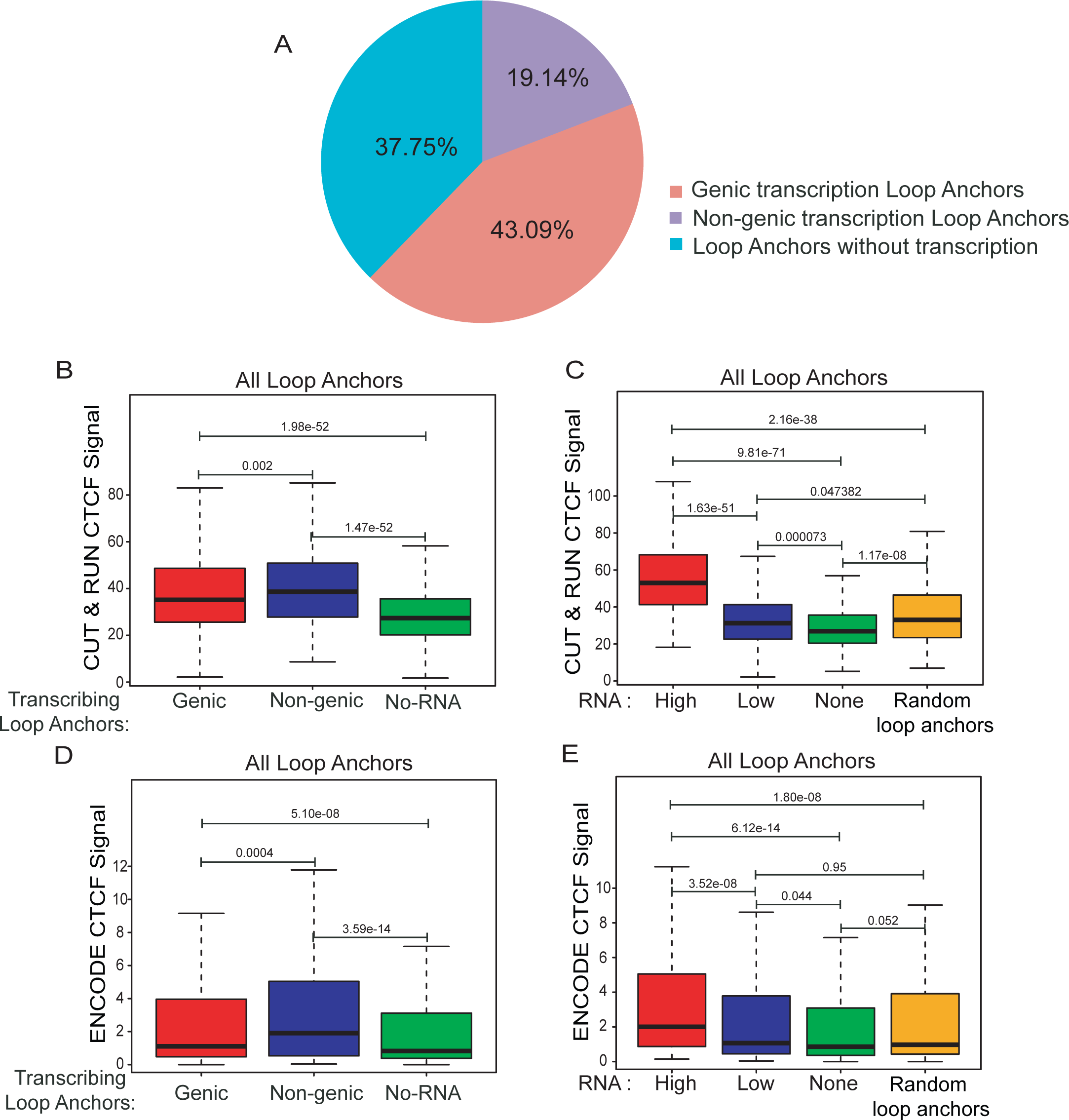
Transcribed Loop Anchors exhibit more CTCF enrichment: **A.** Pie chart showing the percentage of loop anchors that overlap with regions of genic transcription and non-genic de novo transcription and regions without transcription. **B.** CTCF enrichment (CTCF CUT&RUN) at transcribed genic, non-genic and non-transcribed loop anchors. **C.** CTCF enrichment on all transcribed loop anchors with high and low RNA versus non-transcribed and random loop anchors. **D.** CTCF enrichment (CTCF ChIP-seq) at transcribed genic, non-genic and non-transcribed loop anchors. **E.** CTCF enrichment at all transcribed loop anchors with high and low RNA versus non-transcribed and random loop anchors. p-values in boxplots were calculated by Wilcoxon rank-sum test. The boxplots depict the minimum (Q1-1.5*IQR), first quartile, median, third quartile and maximum (Q3+1.5*IQR) without outliers.

**Supp Figure 4.**
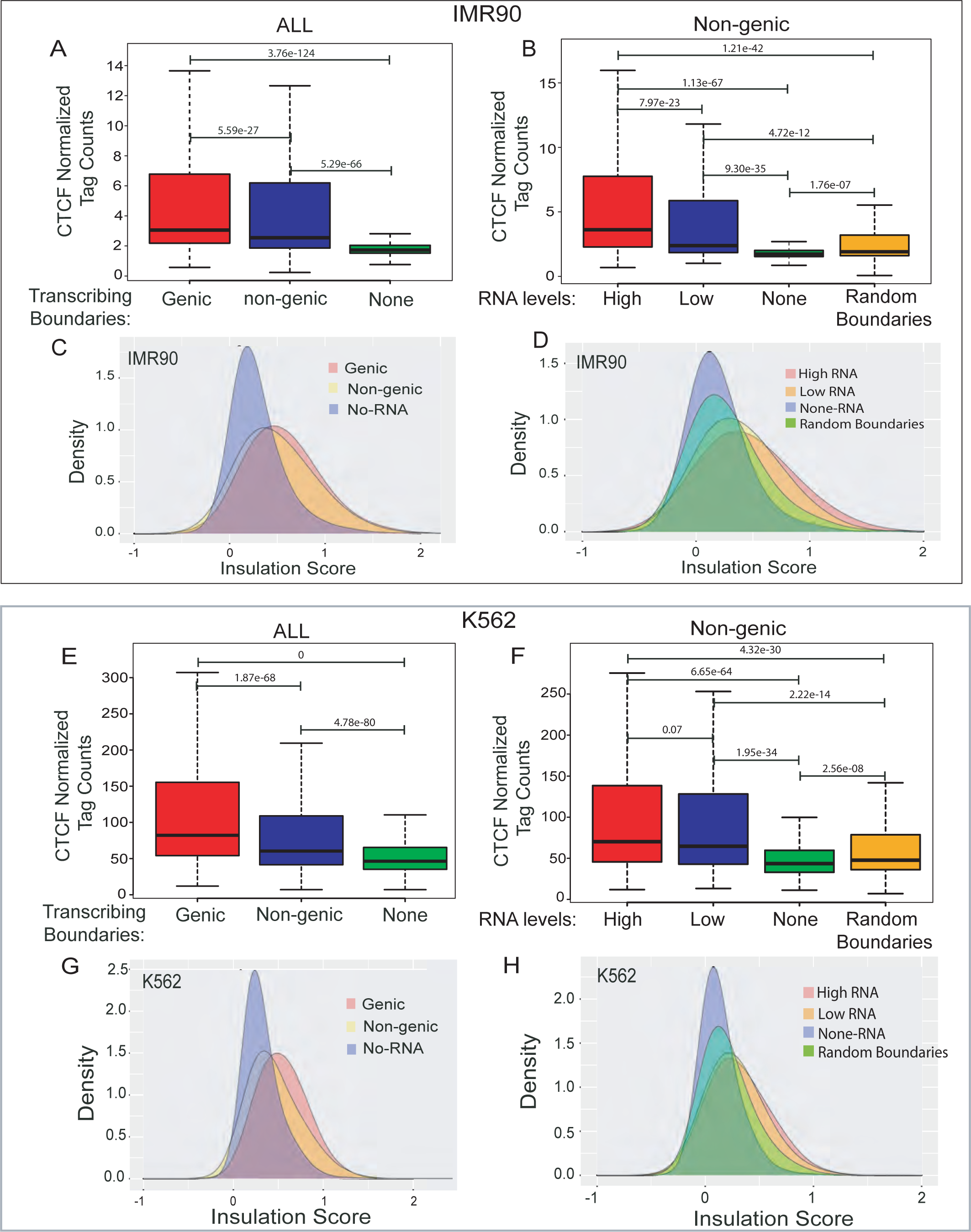
Transcribed boundaries in IMR90 and K562 exhibit better insulation: **A.** CTCF enrichment at transcribed genic, non-genic and non-transcribed boundaries in IMR90 cells. **B.** CTCF enrichment at non-genic transcribed boundaries with high and low non-coding RNA versus non-transcribed and random boundaries in IMR90 cells. **C.** Density plots showing insulation scores for transcribed genic, non-genic and non-transcribed boundaries in IMR90 cells**. D.** Density plots showing insulation scores for non-genic transcribed boundaries exhibiting varying levels of non-coding RNAs versus non-transcribed and random boundaries in IMR90. **E-F**. Same as A-B in K562 cells. **G-H.** Same as C-D in K562 cells. p-values in boxplots were calculated by Wilcoxon rank-sum test. The boxplots depict the minimum (Q1-1.5*IQR), first quartile, median, third quartile and maximum (Q3+1.5*IQR) without outliers.

**Supp Figure 5.**
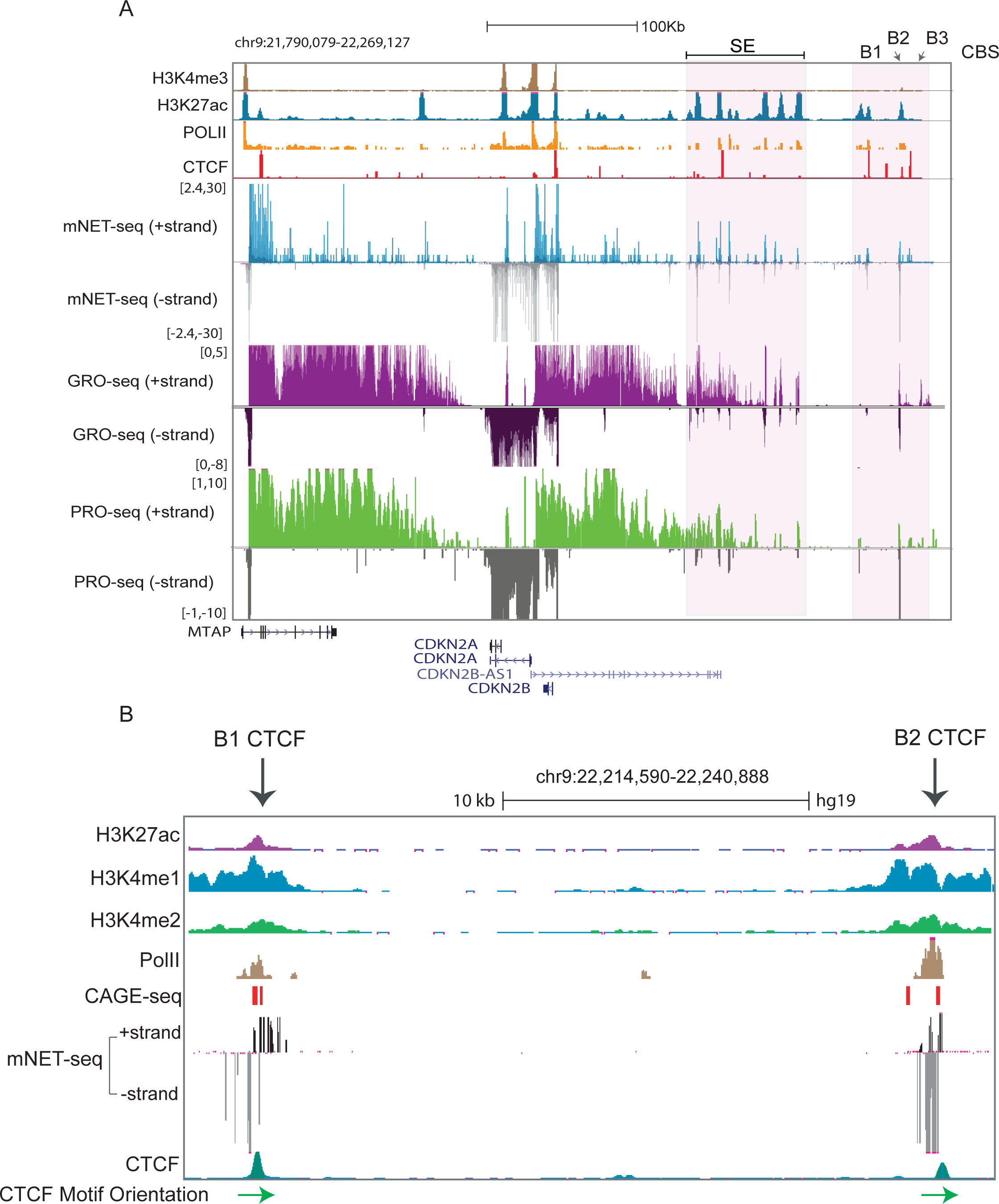
Browser shots of the *INK4a/ARF* loop domain: **A.** A browser shot showing the *INK4a/ARF* locus in the 9p21 region. Below are H3K4me3, H3K27ac (enhancers), PolII, CTCF, mNET-seq tracks, GRO-seq tracks, PRO-seq tracks and gene annotations. The highlighted region shows the B1, B2 and B3 CTCF sites at the 3’ boundary and super-enhancer. **B**. Browser shot showing the zoomed-in region of the 3’ boundary (B1 and B2 CTCF sites). Below are H3K27ac, H3K4me1, H3K4me2, PolII, CAGE seq tracks, mNET-seq tracks and CTCF peaks. Arrowheads at the bottom show CTCF motif orientation at the B1 and B2 CTCF sites.

**Supp Figure 6.**
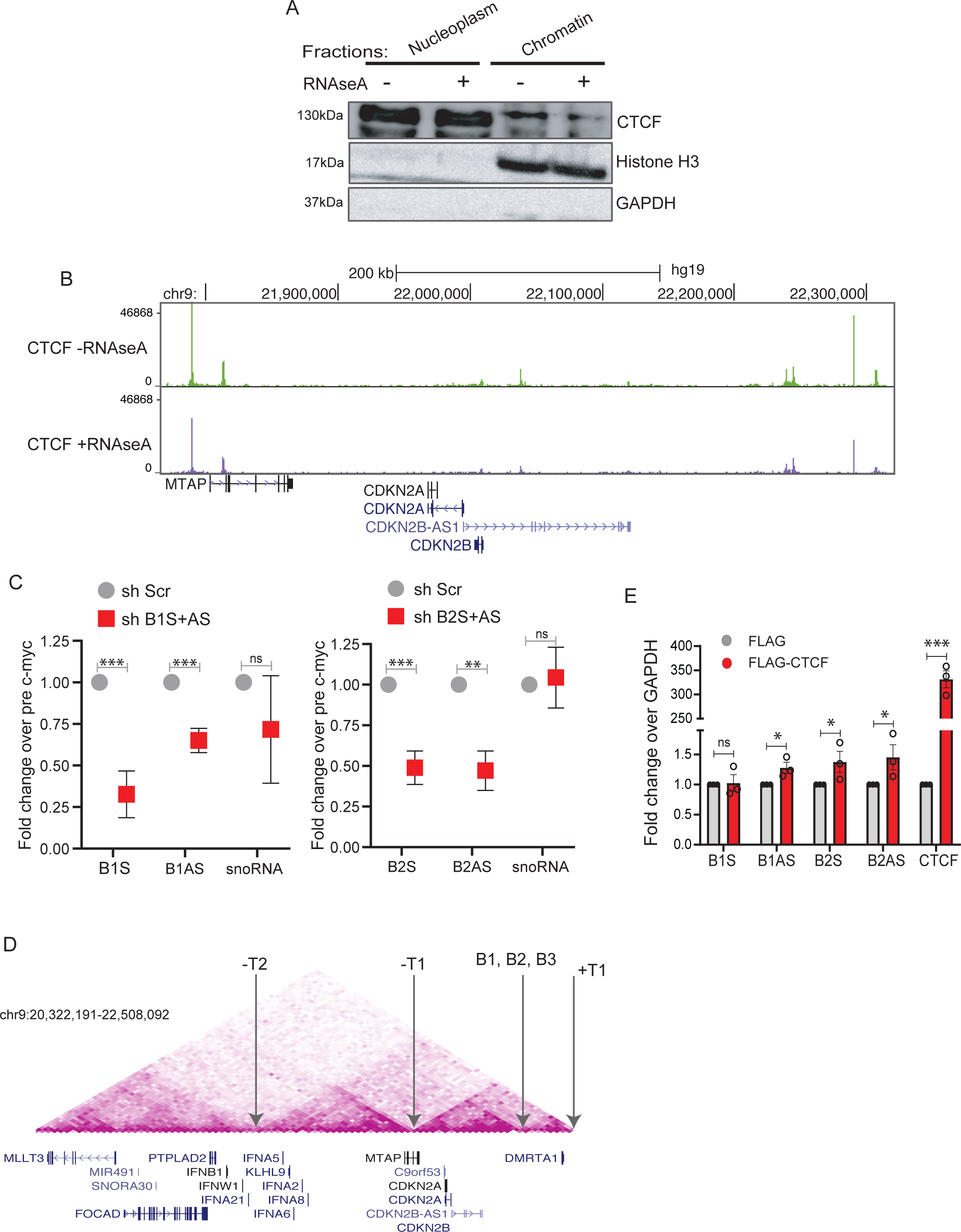
RNA increases CTCF occupancy at TAD boundary: **A.** Immunoblotting with CTCF, Histone H3 and GAPDH on soluble nucleoplasm and chromatin**-**bound fractions of nuclei treated with or without RNaseA. **B.** Browser shot showing CTCF CUT&RUN peaks at the *INK4a/ARF* TAD with and without RNaseA treatment. **C.** qRT-PCRs showing the levels of sense and anti-sense non-coding RNA at B1and B2 CTCF sites and of snoRNA upon B1 and B2 RNA knockdown in the chromatin associated RNA fraction. **D.** TADs around the *INK4a/ARF* TAD. Positions of arrows show the CTCF sites at upstream and downstream TAD boundaries where CTCF occupancy was interrogated. **E.** qRT-PCRs showing the fold-change in sense and anti-sense RNAs from B1 and B2 CTCF sites upon Flag-CTCF overexpression. The last two bars show the increase in CTCF levels. Error bars denote SEM from three biological replicates. p-values were calculated by Student’s two-tailed unpaired t test in **C** and **E**. \*\*\*\**p* < 0.0001, \*\*\**p* < 0.001, \*\**p* < 0.01, \**p* < 0.05, ^ns^p > 0.05.

**Supp Figure 7.**
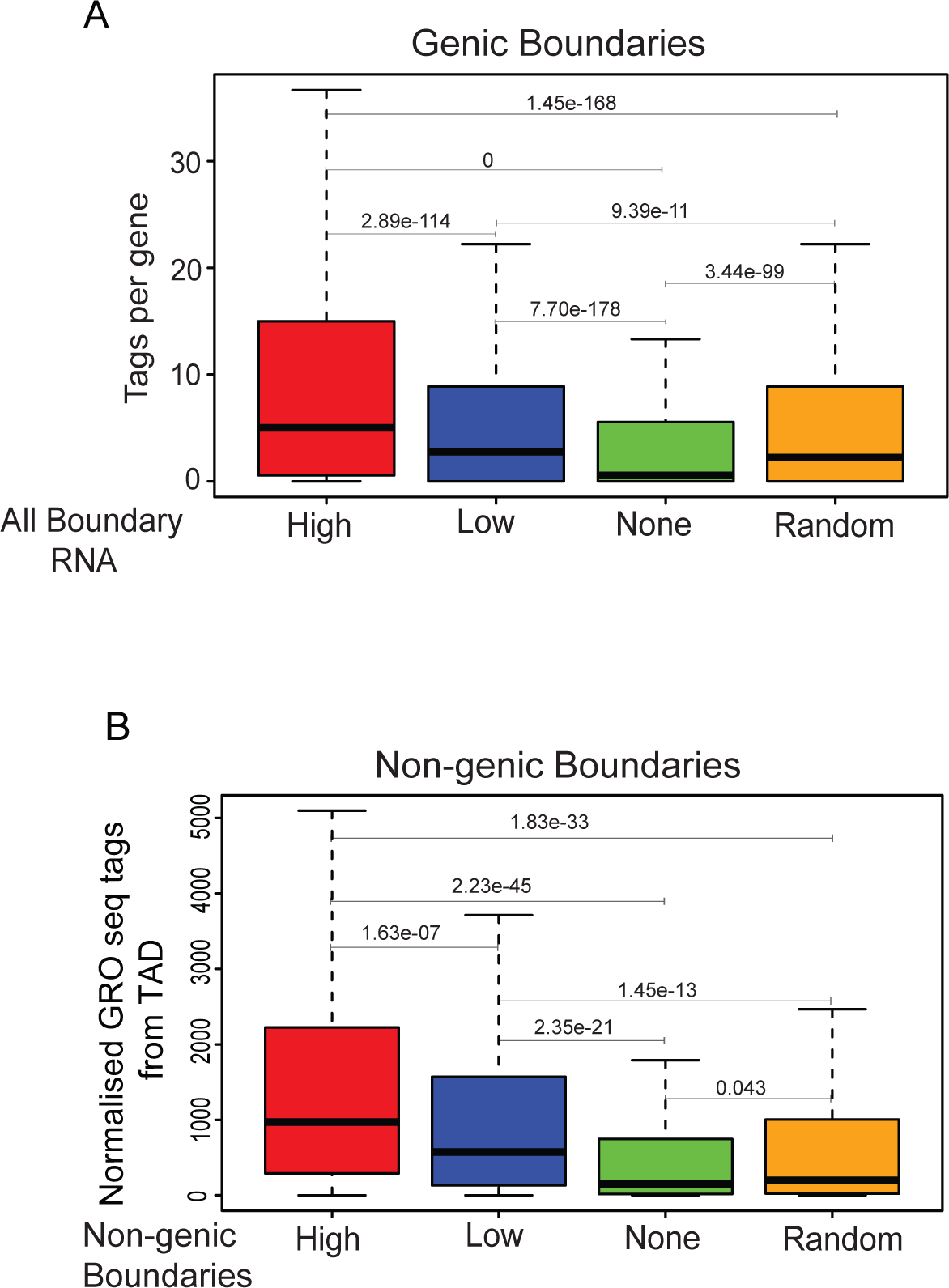
TAD transcription is correlated with transcribed boundaries: **A.** Box plots showing the normalized GRO-seq reads per gene within TADs of high, low, non-transcribed and random boundaries. **B.** Box plots showing the normalized GRO-seq tags within the TADs of transcribed non-genic boundaries with high and low boundary RNAs, non-transcribed and random boundaries.

**Supp Figure 8.**
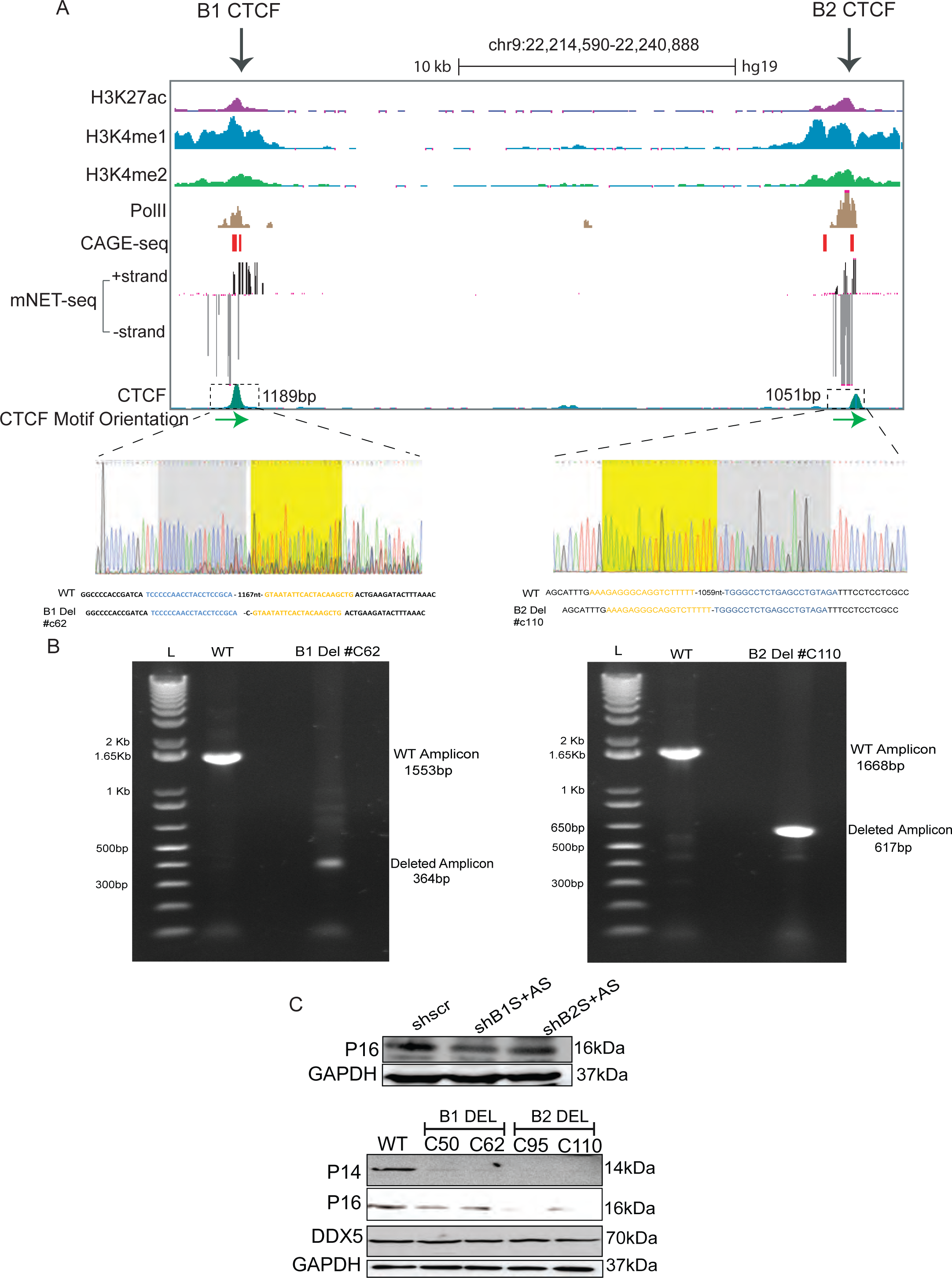
Deletion of B1 and B2 CTCF sites in the *INK4a/ARF* TAD boundary: **A.** UCSC genome browser track depicting the deleted B1 and B2 CTCF sites (boxed region at the bottom). Below are CAGE, NET-seq signal, CTCF peaks and CTCF orientation (Green arrows). The tracks zoom in to the chromatograms from Sanger sequencing on deleted amplicons to verify the deletion. **B.** Agarose gel images showing the amplicons from a Surveyor assay of B1 and B2 deletions. **C.** Immunoblot showing the protein levels of p16 after Scr shRNA knockdown against sense and anti-sense RNA at B1 and B2 CTCF sites (Upper panel). Immunoblot showing the protein levels of p14, p16, DDX5 and GAPDH in WT and in cells with B1 and B2 CTCF site deletions (Lower panel).

**Supp Figure 9.**
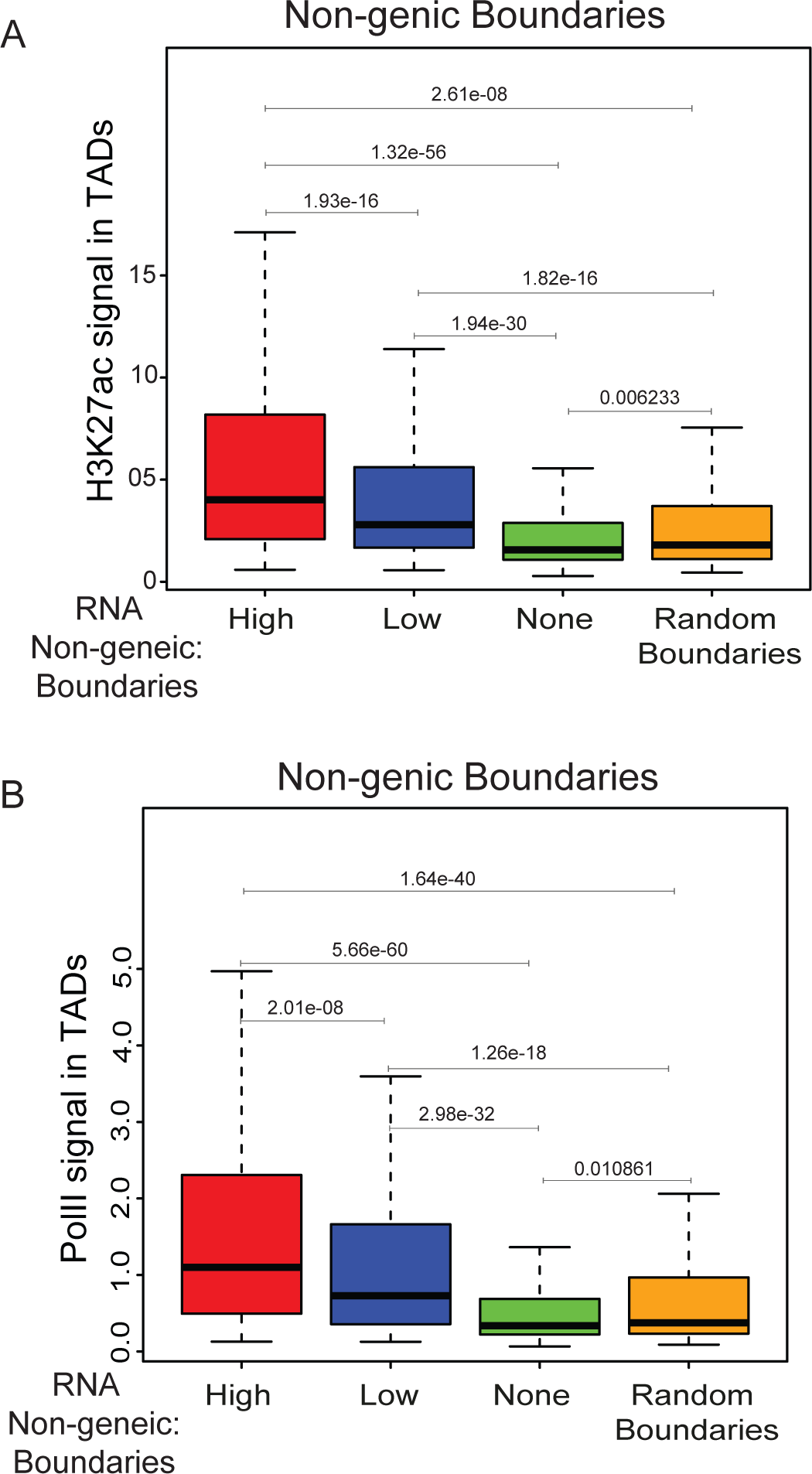
TADs with transcribed boundaries harbor more active enhancers: **A.** Boxplots displaying enrichment of H3K27ac in TADs with high and low transcribed non-genic HeLa boundaries versus non-transcribed and random boundaries. **B.** Levels of PolII in TADs with different non-genic transcribed boundaries vs. non-transcribed and random boundaries.

**Supp Figure 10.**
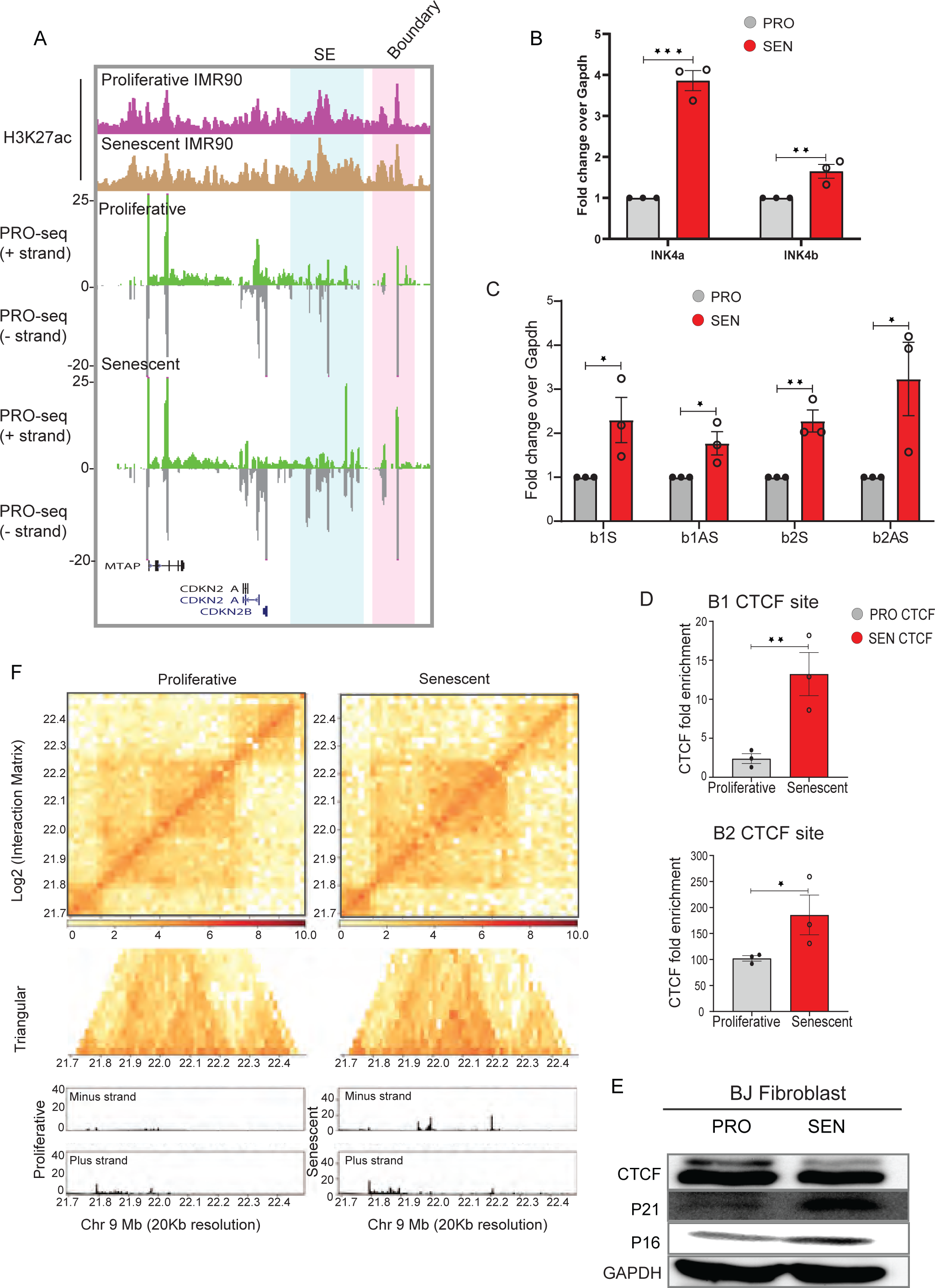
Transcriptional state of *INK4a/ARF* TAD in senescence : **A.** Browser images showing H3K27ac, PRO-seq reads on *INK4a/ARF* TAD from young and replicative senescent IMR90 fibroblasts. Highlighted regions depict the super-enhancer and the boundary. **B**. qRT-PCR plots show the fold change in expression of *INK4a* and *INK4b* in young and DNA damage-induced senescent BJ fibroblasts. **C.** qRT-PCR plots show the fold change in expression of boundary RNA from B1 and B2 CTCF sites in young and DNA damage-induced senescent BJ fibroblasts. **D.** ChIP qRT-PCRs showing the changes in fold enrichment of CTCF at B1 and B2 CTCF sites in young and DNA damage-induced senescent BJ fibroblasts. Error bars denote SEM from three biological replicates. *p*-values were calculated by Student’s two-tailed unpaired *t*-test in **B**, **C**, and **D.** \*\*\**p* < 0.001, \*\**p* < 0.01, \**p* < 0.05, ^ns^*p* > 0.05. **E.** Immunoblot showing the changes in protein levels of CTCF, p21, p16 and GAPDH in young and DNA damage-induced senescent BJ fibroblasts. **F.** *INK4a/ARF* TAD structure in proliferating and senescent cells. Lower boxes represent the transcriptional levels.

**Table 1:**
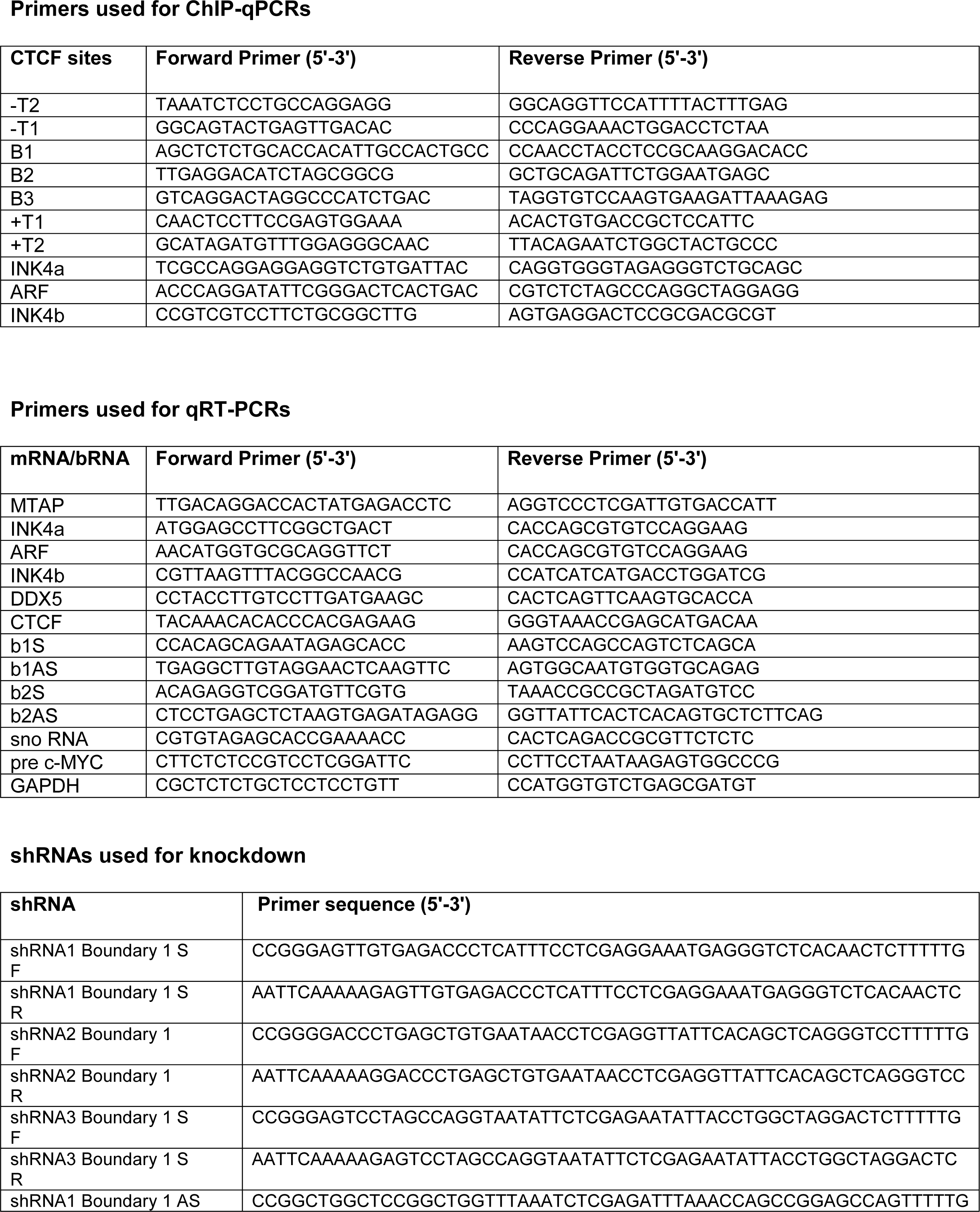
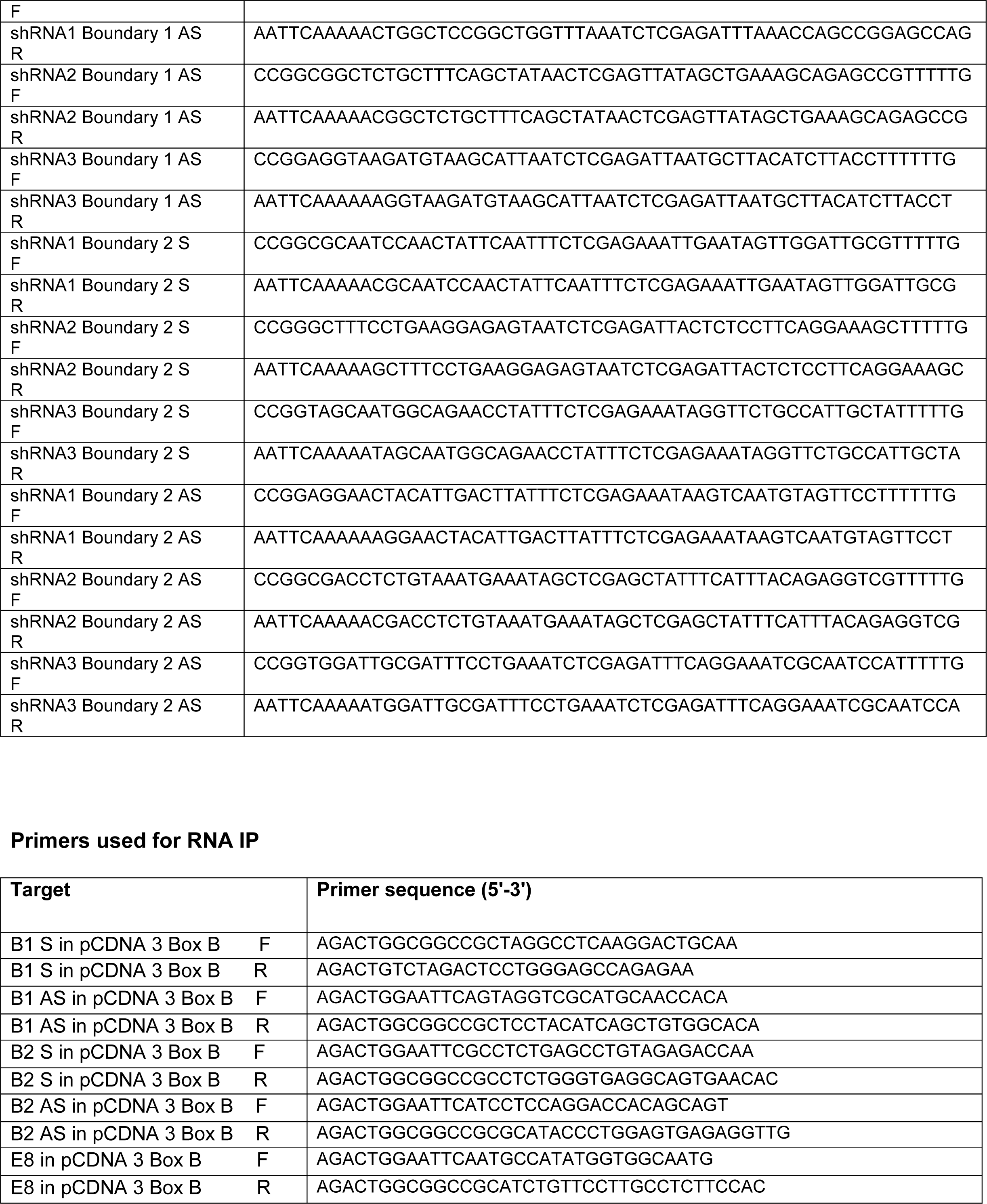
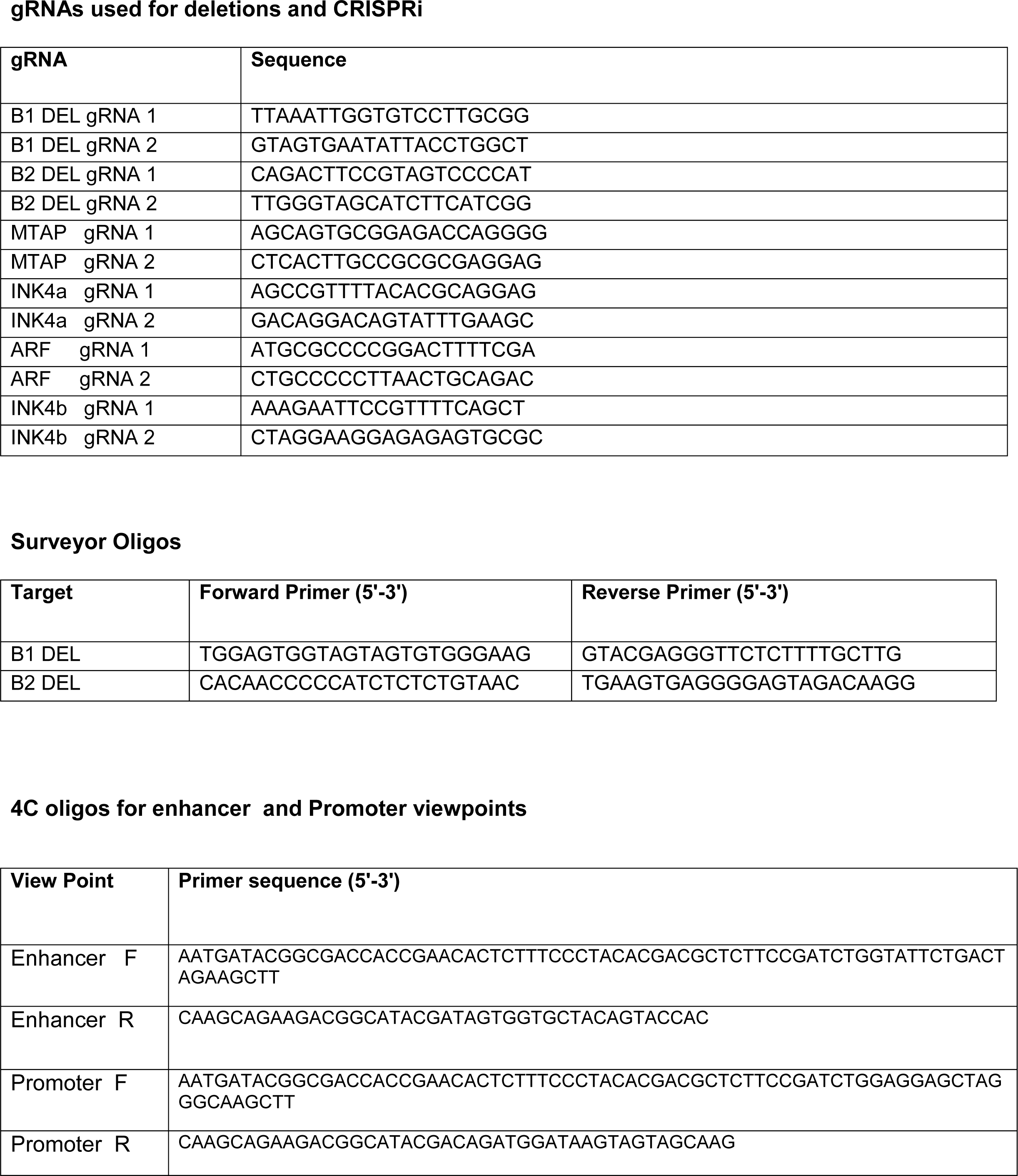
Oligonucleotide sequences used in the study.

**Table 3:**
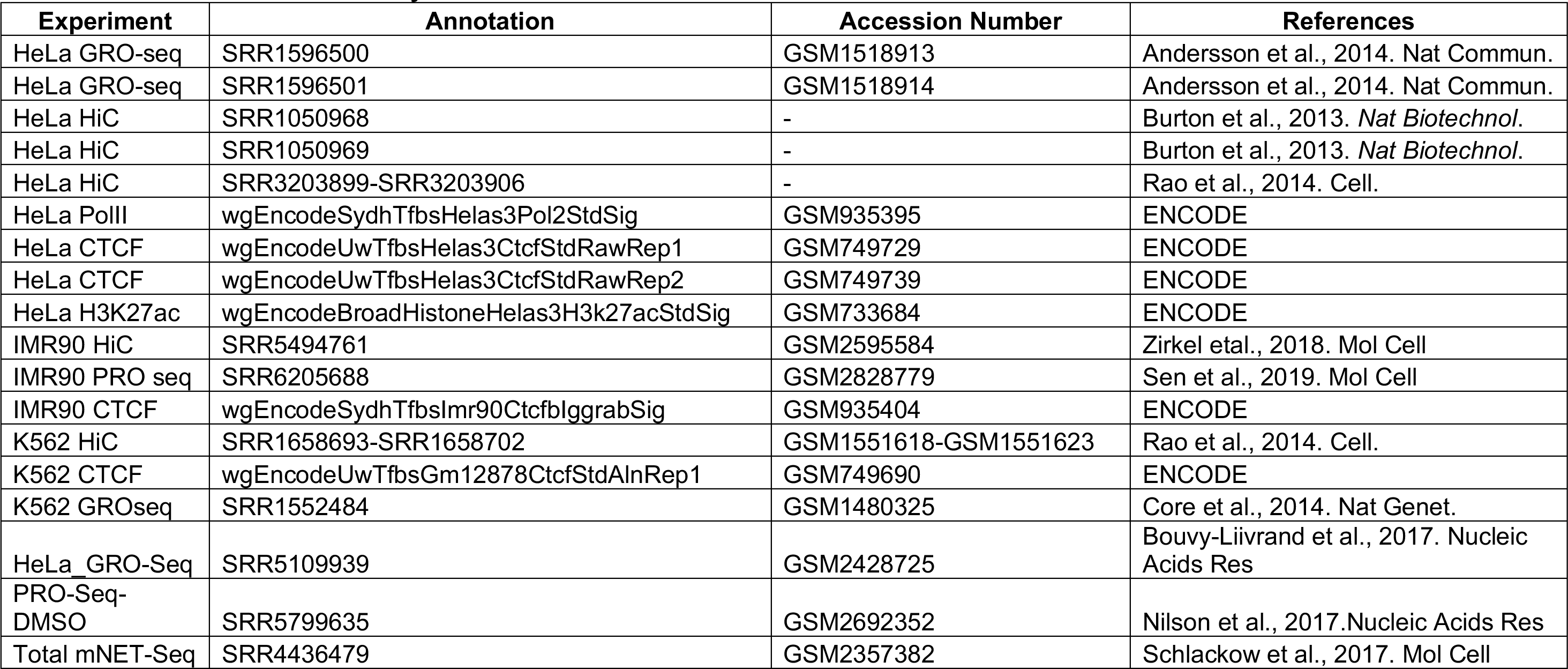
Datasets used in the study.

